# Ribosomal proteins could explain the phylogeny of *Bacillus* species

**DOI:** 10.1101/2020.07.25.221481

**Authors:** Wenfa Ng

## Abstract

Protein translation is a highly conserved process in biology. As participants of translation, ribosomal proteins in the large and small subunits of the ribosomes are likely to be highly conserved; thus, could they be endowed with sufficient sequence diversity to chronicle the evolutionary history of different species in the same or different genus? Using different *Bacillus* species as a model system, this study sought to examine if ribosomal proteins could reproduce the maximum likelihood phylogeny described by 16S rRNA of the investigated *Bacillus* species. *Bacillus* species investigated were *Bacillus amyloliquefaciens, Bacillus cereus, Bacillus licheniformis, Bacillus megaterium, Bacillus pumilus, Bacillus subtilis*, and *Bacillus thuringiensis*. Results revealed that ribosomal proteins could be categorized into four different groups depending on their extent in reproducing the 16S rRNA phylogeny of the different *Bacillus* species. The first group comprises ribosomal protein that could reproduce all the phylogenetic positions of the *Bacillus* species accurately. These ribosomal proteins were ribosomal protein L6, L7/12, L9, L13, L24, L32, S3, S9, S12, S15, S16, S17, and S18. Ribosomal proteins that hold partial phylogenetic significance constitutes the second group where the ribosomal proteins could reproduce the major branches of the 16S rRNA phylogenetic tree but differ in the placement of one or two *Bacillus* species. In general, this group of ribosomal proteins had difficulty differentiating *B. licheniformis* and *B. pumilus* at the sequence level. Members of this group of ribosomal protein include ribosomal protein L22, L29, L30, L31 Type B, L33, L35, S1, S4, S5, S6, S7, S8, S11, S13, S19, and S20. The third group of ribosomal proteins were those which were highly conserved at the sequence level, and which could not differentiate the different *Bacillus* species. These ribosomal proteins were ribosomal protein L5, L36, S2, S10, and S21. Finally, there were also ribosomal proteins that randomly placed the different *Bacillus* species into phylogenetic positions not in sync with those depicted by the 16S rRNA phylogenetic tree. These ribosomal proteins were ribosomal protein L7Ae, L17, L20, L23, L27, L28, L31, L34 and S14 Type Z. Overall, members of all four groups of ribosomal proteins came from both the large and small ribosome subunits which meant that evolutionary forces exerted selective pressure on both subunits but at differing extents. Collectively, specific ribosomal proteins could reproduce the phylogeny of different *Bacillus* species as described by the gold standard phylogenetic marker, 16S rRNA, which highlighted that co-evolutionary processes could be at work in shaping the evolution of ribosomal proteins and rRNA in close contact with each other in the ribosome.

**Subject areas:** ecology, biochemistry, biotechnology, microbiology, cell biology

**Significance of the work:** 16S rRNA is the gold standard phylogenetic marker used to inform the evolutionary relationships between different species across the three domains of life. Given that 16S rRNA is nestled in the ribosomes together with a consortium of ribosomal proteins each with unique structural and enzymatic functions, could ribosomal proteins be used similarly as phylogenetic markers for informing species divergence and relationships? Specifically, as part of the highly conserved ribosome important to protein translation, do ribosomal proteins possess sufficient sequence diversity to help chronicle the evolutionary relationships between different species? By reconstructing the maximum likelihood phylogenetic tree of different *Bacillus* species, this study revealed that ribosomal proteins fall into four categories concerning their utility for informing phylogeny between different species of the same genus. Specifically, there existed ribosomal proteins able to accurately reproduce the phylogenetic tree described by 16S rRNA. On the other hand, there were ribosomal proteins that hold only partial phylogenetic significance where they could reproduce the major branches of the reference phylogenetic tree but differ in the placement of one or two species along the tree. Besides the above two categories, they were also ribosomal proteins whose sequence diversity was not sufficient to help differentiate between different *Bacillus* species. Finally, another class of ribosomal proteins did not chronicle the evolutionary trajectories of the different species resulting in phylogenetic tree with random placement of the different species. Overall, evolutionary forces likely exerted different selection forces on different ribosome subunits as well as individual ribosomal protein that resulted in the differentiation of their utility as phylogenetic markers of different species of the same genus. Co-evolution between ribosomal proteins as well as between ribosomal proteins and rRNA might underpin part of the evolutionary history chronicled by individual ribosomal proteins, thereby, endowing them with phylogenetic significance.

**Highlights:**

1. Ribosomal proteins were found to be useful in describing the phylogeny of different *Bacillus* species compared to the gold standard phylogenetic marker, 16S rRNA.
2. By examining the maximum likelihood phylogenetic tree reconstructed, ribosomal proteins could be categorized into four groups with differing phylogenetic significance.
3. The first group comprises ribosomal proteins able to accurately reproduce all the phylogenetic positions of different *Bacillus* species relative to 16S rRNA phylogenetic tree. This group include ribosomal protein L6, L7/12, L9, L13, L24, L32, S3, S9, S12, S15, S16, S17, and S18.
4. The second group refers to ribosomal proteins able to reproduce the major branches of the 16S rRNA phylogenetic tree but lacks in the correct placement of one or two *Bacillus* species. These ribosomal proteins were L22, L29, L30, L31 Type B, L33, L35, SI, S4, S5, S6, S7, S8, Sll, S13, S19, and S20.
5. The third group of ribosomal proteins are ones with highly conserved sequence unable to differentiate between different *Bacillus* species. It comprised ribosomal proteins L5, L36, S2, S10, and S21.
6. The final group of ribosomal proteins did not chronicle the evolutionary forces acting on the different *Bacillus* species and generated phylogenetic trees with random placement of the different *Bacillus* species. These ribosomal proteins were L7Ae, L17, L20, L23, L27, L28, L31, L34 and S14 Type Z.

## Introduction

Phylogenetic relationships between species is typically described by 16S rRNA given the presence of hypervariable regions in the gene that could chronicle evolutionary events that chance on the species.^1,2^ Thus, 16S rRNA owes its ability at describing the evolutionary relatedness of different species to its sequence diversity. Specifically, 16S rRNA is at the sweet spot where it is sufficiently important to overall ribosome function to be conserved in sequence, and yet remains diversified in sequence at hypervariable regions. This indicated that 16S rRNA might not be absolutely critical to ribosome function, where otherwise it would have been much more highly conserved and lacks sequence diversity necessary to chronicle evolutionary events.

Thus, to be useful for taxonomic purposes, a biomolecule must be sufficiently important to cellular function to be conserved in sequence and structure, and yet possess sufficient sequence space for cataloguing the myriad evolutionary pressure that acts on a species. Ribosomal proteins that help constitute the ribosome large and small subunit in structure and function seems suitable candidates for exploring their phylogenetic potential. Specifically, nestled within the structural space of the two ribosome subunits, ribosomal proteins could have undergone co-evolution with 16S rRNA; for example, through the need to maintain particular structural fold or function within and between the ribosome subunits.

Efforts aimed at understanding the phylogenetic potential of ribosomal proteins came about partially through attempts to annotate the mass spectra of microbial species profiled by the gentle ionisation technique of matrix-assisted laser desorption/ionization time-of-flight mass spectrometry (MALDI-TOF MS).^3,4^ Known to be a relatively new technique capable of identifying microorganisms down to the species and strain level,^5,6^ MALDI-TOF MS microbial identification has long suffered from the inability to annotate many of the mass peaks profiled from microbial species. Given the essential nature of ribosomal proteins to cellular function, efforts to annotate ribosomal protein mass peaks in MALDI-TOF mass spectra of bacterial species informed of the relative importance of the class of proteins to potentially inform the phylogeny of different bacterial species.^7,8^ Denoted as biomarker proteins in MALDI-TOF mass spectra, ribosomal proteins have been intensively profiled as candidates of mass peaks and for informing species identities.^7-9^ However, only a subset of ribosomal proteins have been annotated as ribosomal protein mass peaks in MALDI-TOF mass spectra;^3^ thus, leaving a significant gap in understanding the potential phylogenetic significance of the whole ensemble of ribosomal proteins of a cell.

Thus, the objective of this study is to understand the phylogenetic potential of individual ribosomal proteins from the ensemble of proteins that constitute the ribosome. Previous studies have gleaned evidence supporting co-evolution between ribosomal RNA and proteins, and more importantly, the ability of ribosomal proteins in defining major phylogenetic lineages.^10^ Given that *Bacillus* species have been known to present difficulties to phylogenetic classification based on MALDI-TOF mass spectra,^11^ this study would reconstruct the phylogenetic tree of seven *Bacillus* species using sequence information from individual ribosomal proteins to help answer the question of whether ribosomal proteins possess sufficient phylogenetic power to differentiate bacteria down to the species level in a hard to differentiate species complex such as *Bacillus* spp. These phylogenetic trees would be assessed for concordance through comparison with the phylogenetic tree reconstructed based on the 16S rRNA sequence information of the *Bacillus* species under study.

## Materials and Methods

*Bacillus* species investigated in this study were *Bacillus amyloliquefaciens, Bacillus cereus, Bacillus licheniformis, Bacillus megaterium, Bacillus pumilus, Bacillus subtilis*, and *Bacillus thuringiensis*. 16S rRNA sequence information for the bacterial species were downloaded from the SILVA database^12^ (https://www.arb-silva.de/). Proteomes of the bacterial species were downloaded from UniProt and the ribosomal proteins of the species extracted from the proteome information. Both 16S rRNA and ribosomal proteins amino acid sequence of the 7 *Bacillus* species were aligned using ClustalW algorithm with default parameters in MEGA 7 software. The aligned nucleotide and amino acid sequences of the 7 *Bacillus* species were subsequently used in reconstruction of maximum likelihood phylogenetic trees using default parameters in MEGA 7^13^ (https://www.megasoftware.net/).

## Results and Discussion

Given that 16S rRNA is the gold standard approach for defining the phylogenetic relationships between species, a maximum likelihood phylogenetic tree based on the 16S rRNA gene sequence of the 7 *Bacillus* species were reconstructed (Figure 1). By comparing the reconstructed phylogenetic trees of individual ribosomal proteins with that based on 16S rRNA, it would be possible to determine if the ribosomal proteins could explain the phylogeny of the *Bacillus* species.

**Figure 1:**
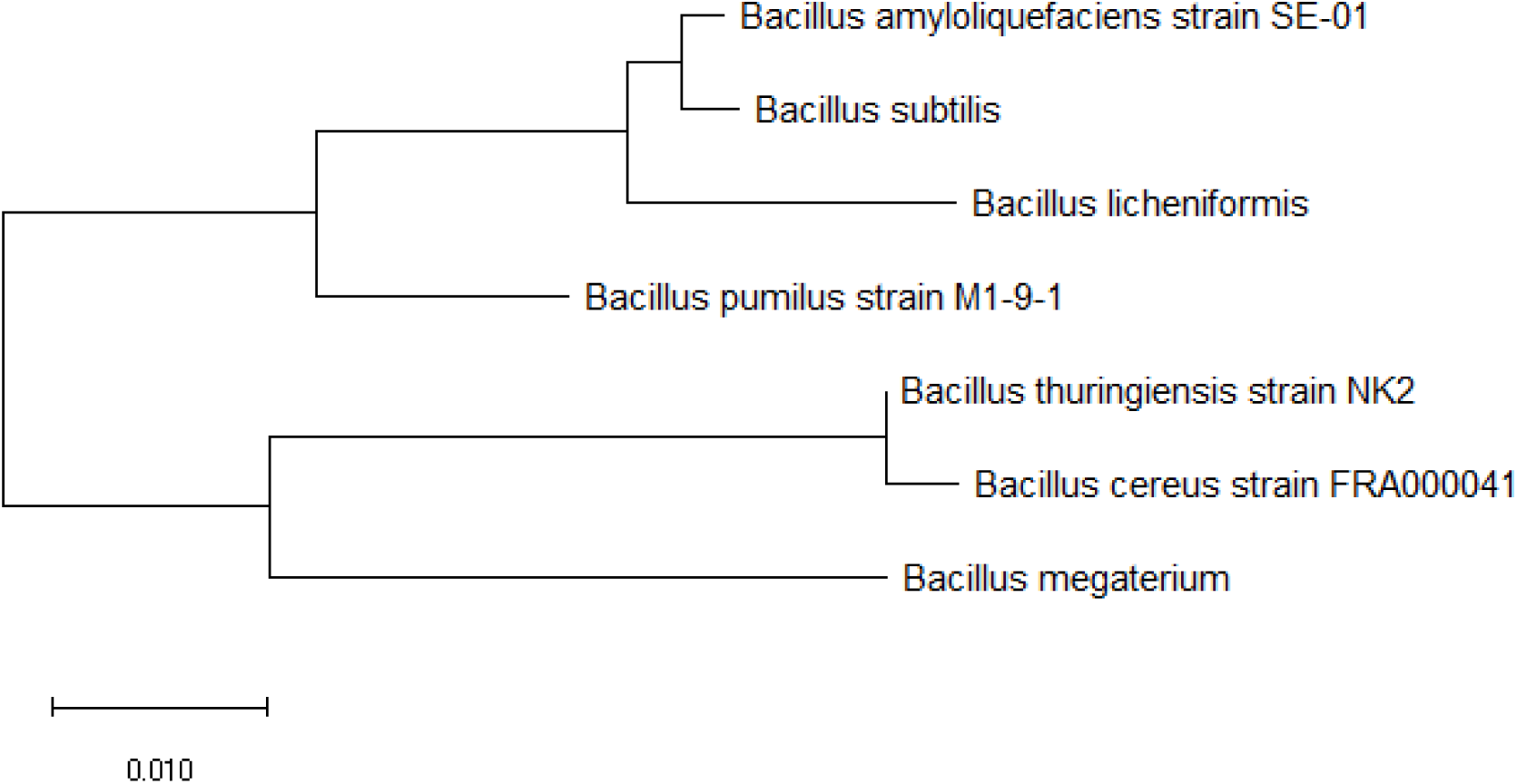
Maximum likelihood phylogenetic tree based on 16S rRNA gene of different *Bacillus* species.

Figure 2 shows the maximum likelihood phylogenetic tree based on ribosomal protein L6 of different *Bacillus* species. The data revealed that ribosomal protein L6 could reproduce the phylogenetic relationships depicted in the phylogenetic tree based on 16S rRNA and thus, the ribosomal protein could explain the phylogeny of the different investigated *Bacillus* species.

**Figure 2:**
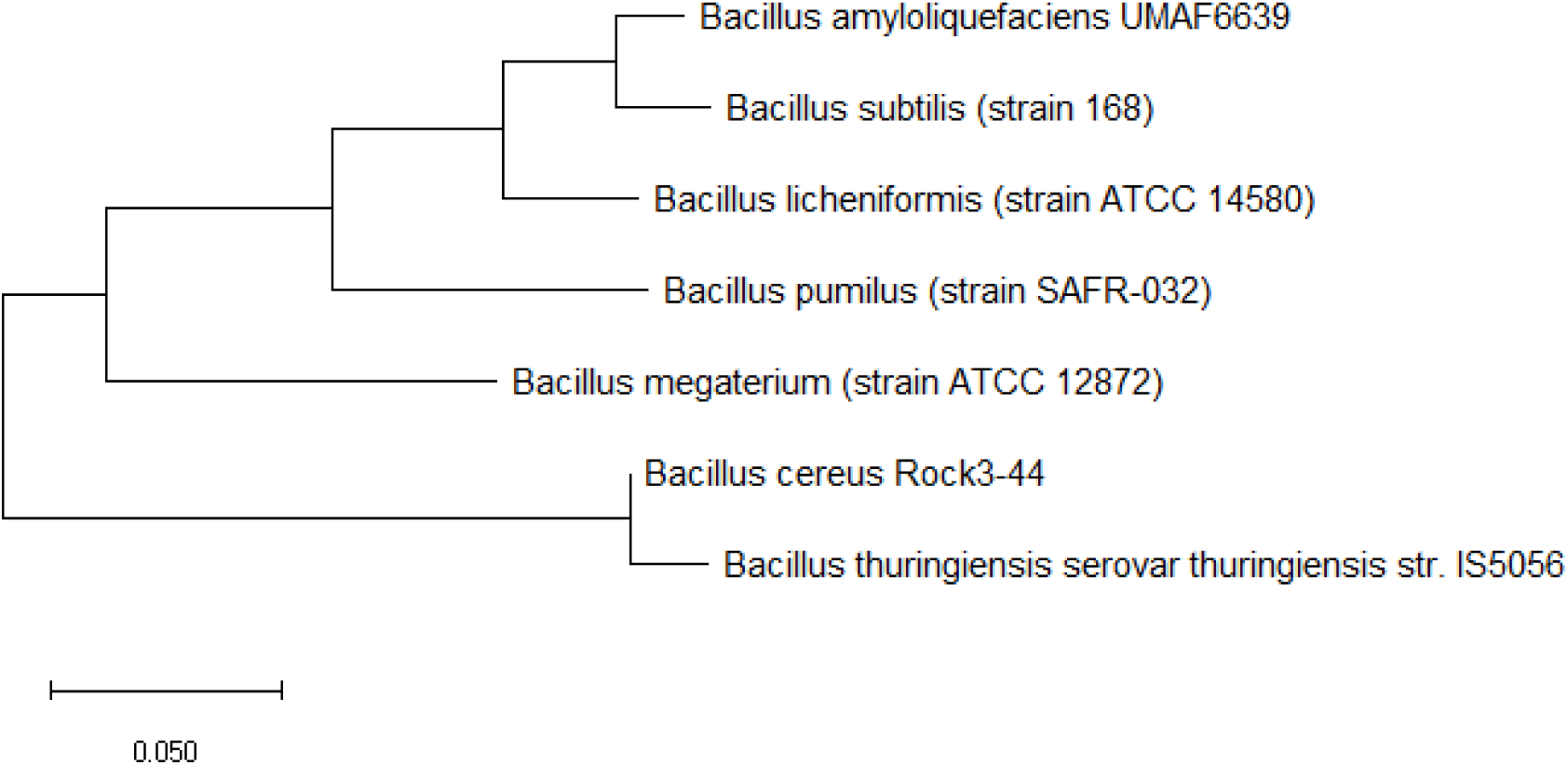
Maximum likelihood phylogenetic tree based on ribosomal protein L6 of different *Bacillus* species.

Figure 3 shows the maximum likelihood phylogenetic tree based on ribosomal protein L7/L12 of different *Bacillus* species. The data revealed that ribosomal protein L7/L12 could reproduce the phylogenetic relationships between different *Bacillus* species as presented by the molecular sequence of 16S rRNA. Thus, ribosomal protein L7/L12 hold phylogenetic significance for explaining the phylogeny between different *Bacillus* species.

**Figure 3:**
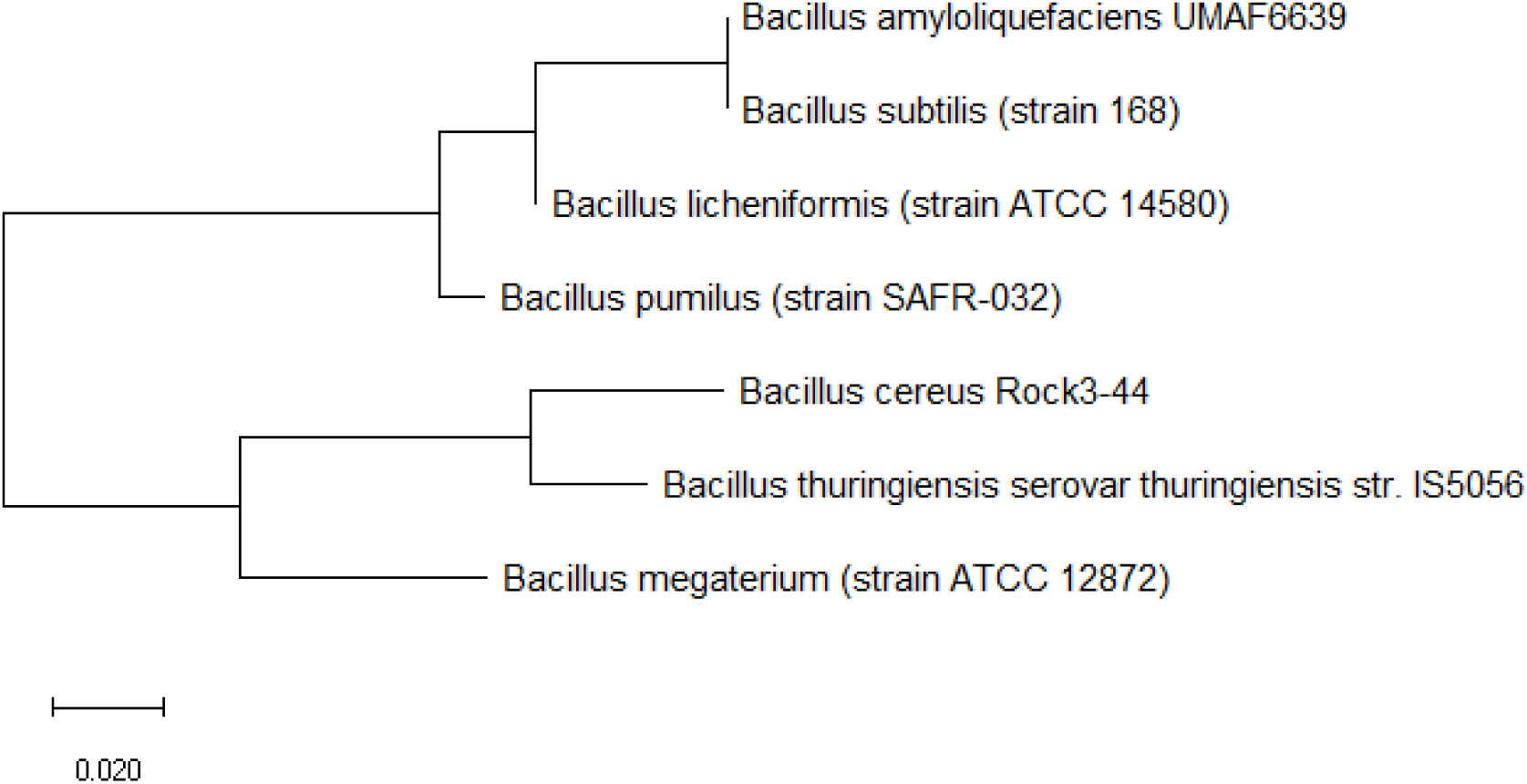
Maximum likelihood phylogenetic tree based on ribosomal protein L7/L12 of different *Bacillus* species.

Figure 4 shows the maximum likelihood phylogenetic tree based on ribosomal protein L9 of different *Bacillus* species. The data revealed that ribosomal protein L9 could reproduce the phylogenetic tree of 16S rRNA for the same *Bacillus* species. Thus, ribosomal protein L9 holds phylogenetic significance in describing the evolutionary relatedness of the different *Bacillus* species.

**Figure 4:**
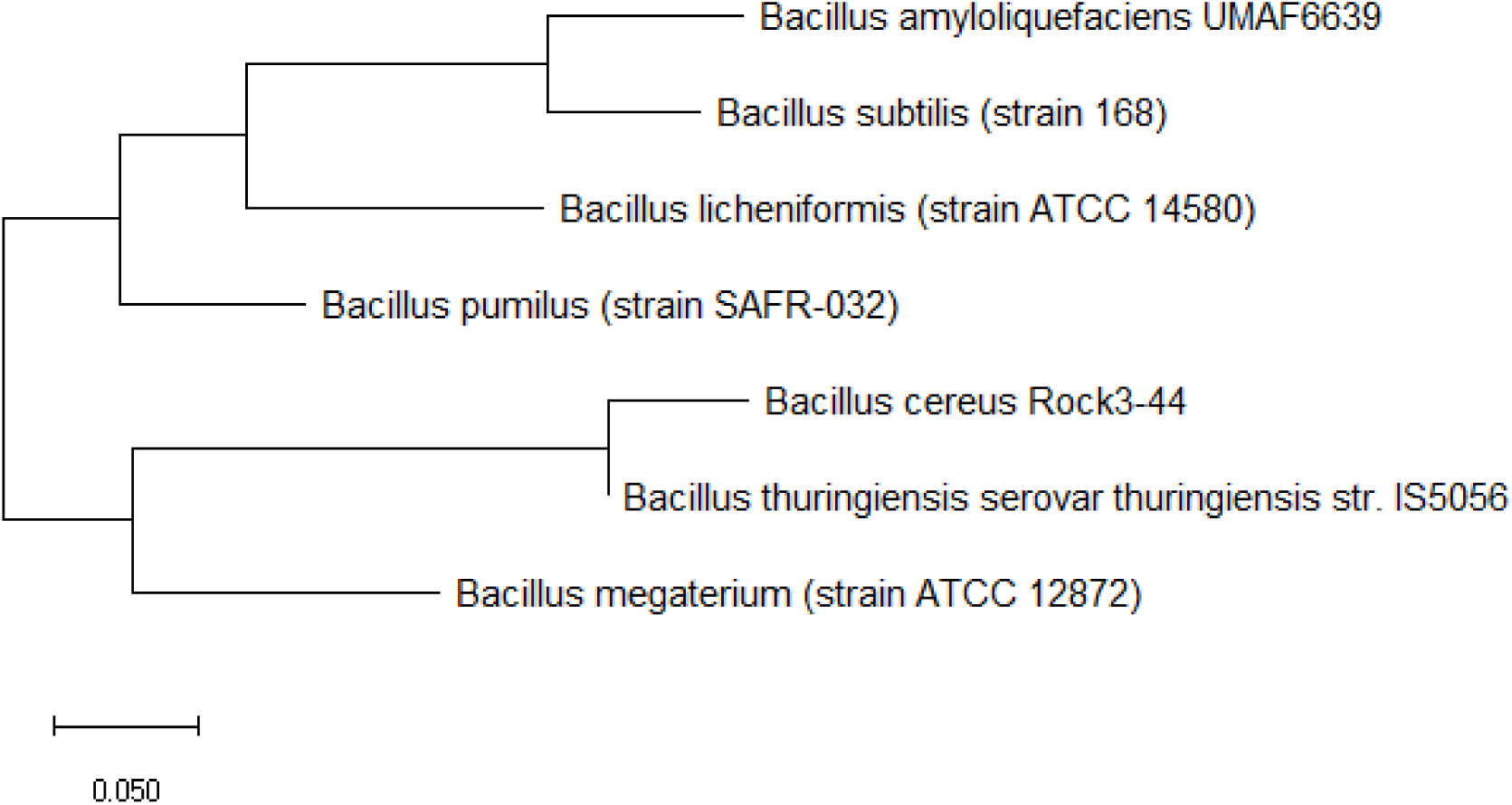
Maximum likelihood phylogenetic tree based on ribosomal protein L9 of different *Bacillus* species.

Maximum likelihood phylogenetic tree based on ribosomal protein L13 of different *Bacillus* species is shown in Figure 5. Specifically, the data revealed that ribosomal protein L13 could reproduce all the branches of the 16S rRNA phylogenetic tree for the same species. Thus, ribosomal protein L13 hold phylogenetic significance for explaining the evolutionary trajectories of the investigated *Bacillus* species.

**Figure 5:**
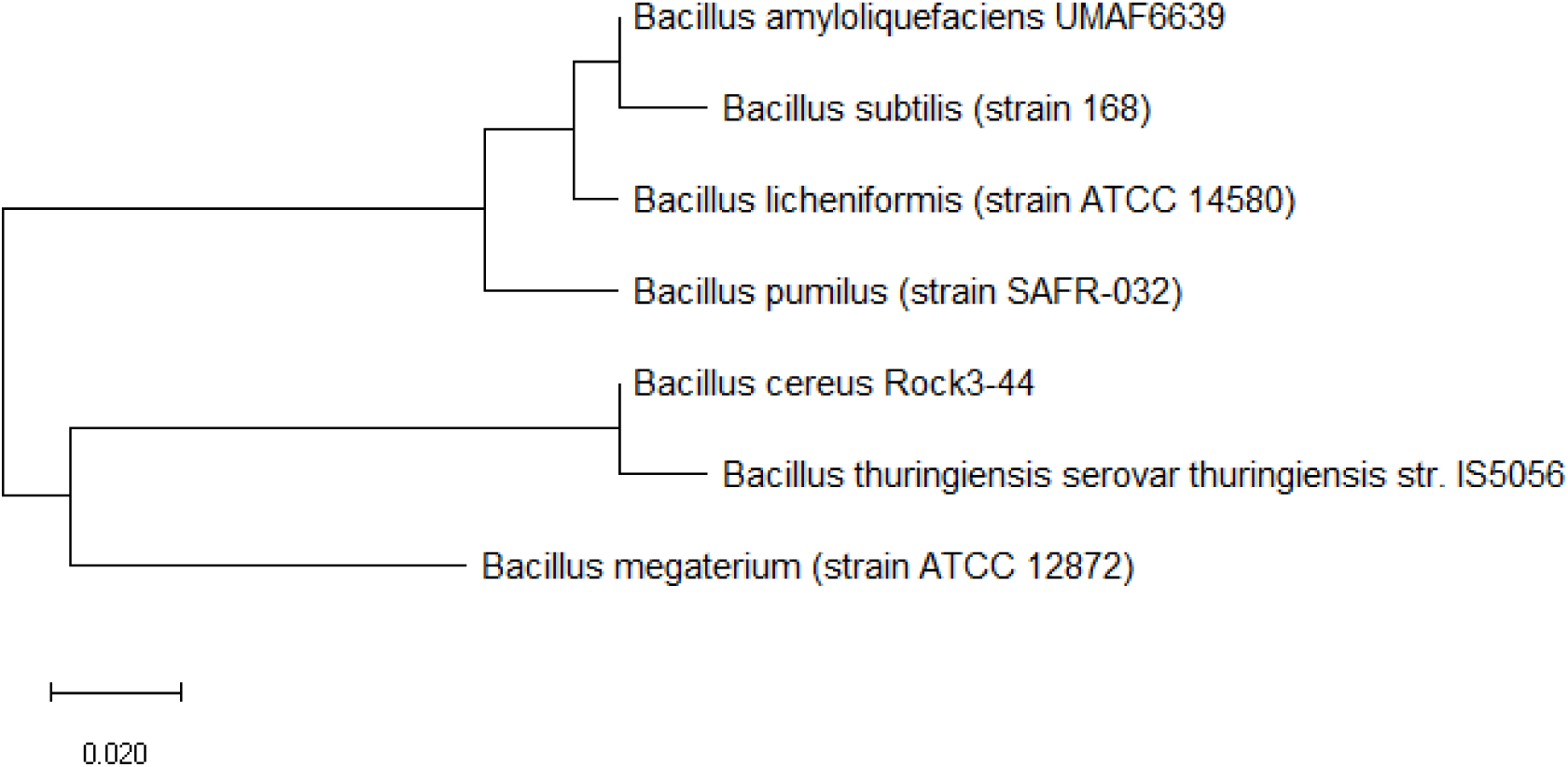
Maximum likelihood phylogenetic tree based on ribosomal protein L13 of different *Bacillus* species.

Figure 6 shows the maximum likelihood phylogenetic tree based on ribosomal protein L18 of different *Bacillus* species. The data revealed that ribosomal protein L18 could reproduce the phylogenetic tree based on 16S rRNA of the same set of *Bacillus* species. Thus, ribosomal protein L18 hold phylogenetic significance for chronicling the evolutionary relationships of the *Bacillus* species.

**Figure 6:**
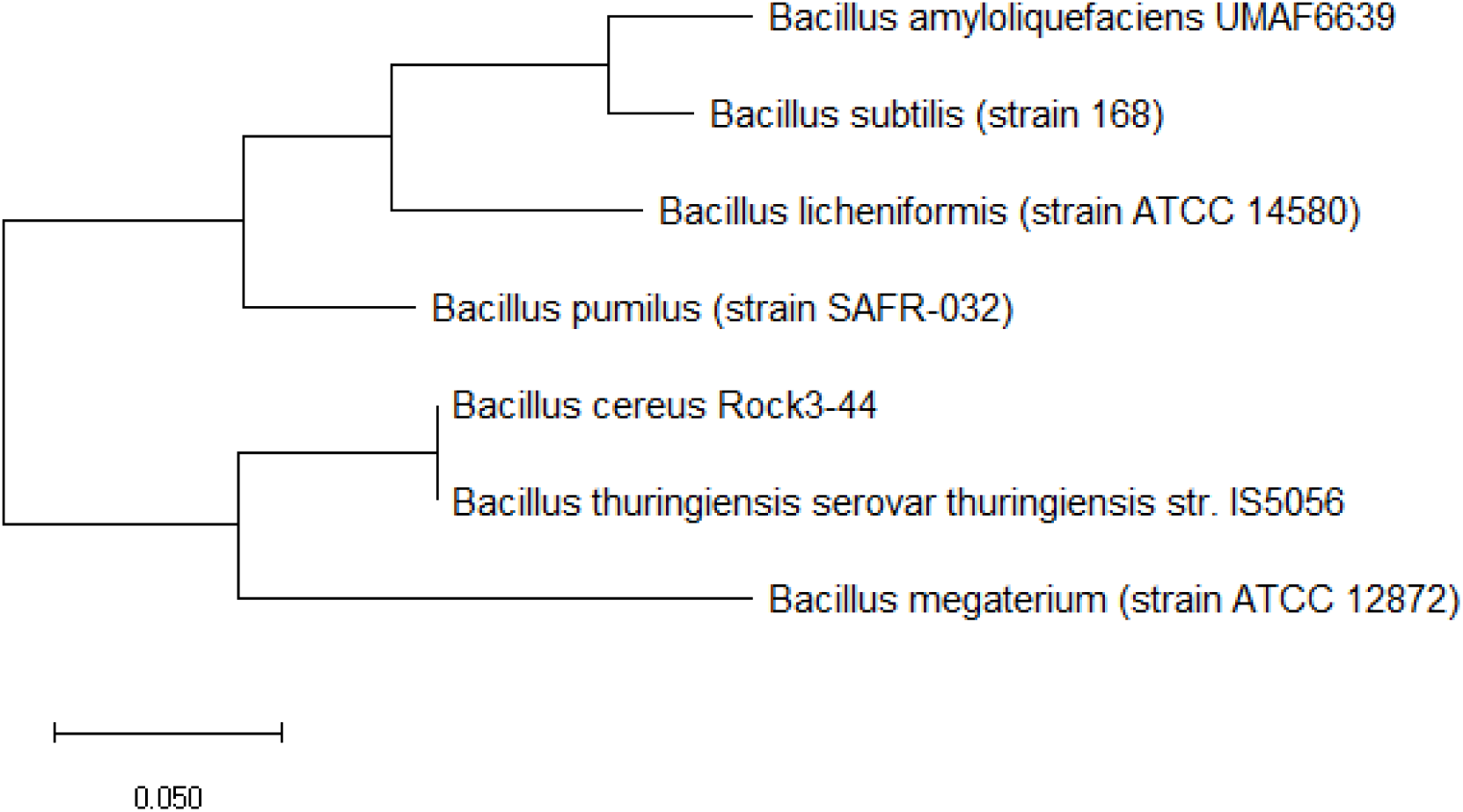
Maximum likelihood phylogenetic tree based on ribosomal protein L18 of different *Bacillus* species.

Figure 7 shows the maximum likelihood phylogenetic tree based on ribosomal protein L19 of different *Bacillus* species. The data revealed that ribosomal protein L19 could reproduce all the major branches of the 16S rRNA phylogenetic tree of the same *Bacillus* species. Thus, ribosomal protein L19 hold phylogenetic significance for explaining the phylogeny of the different *Bacillus* species.

**Figure 7:**
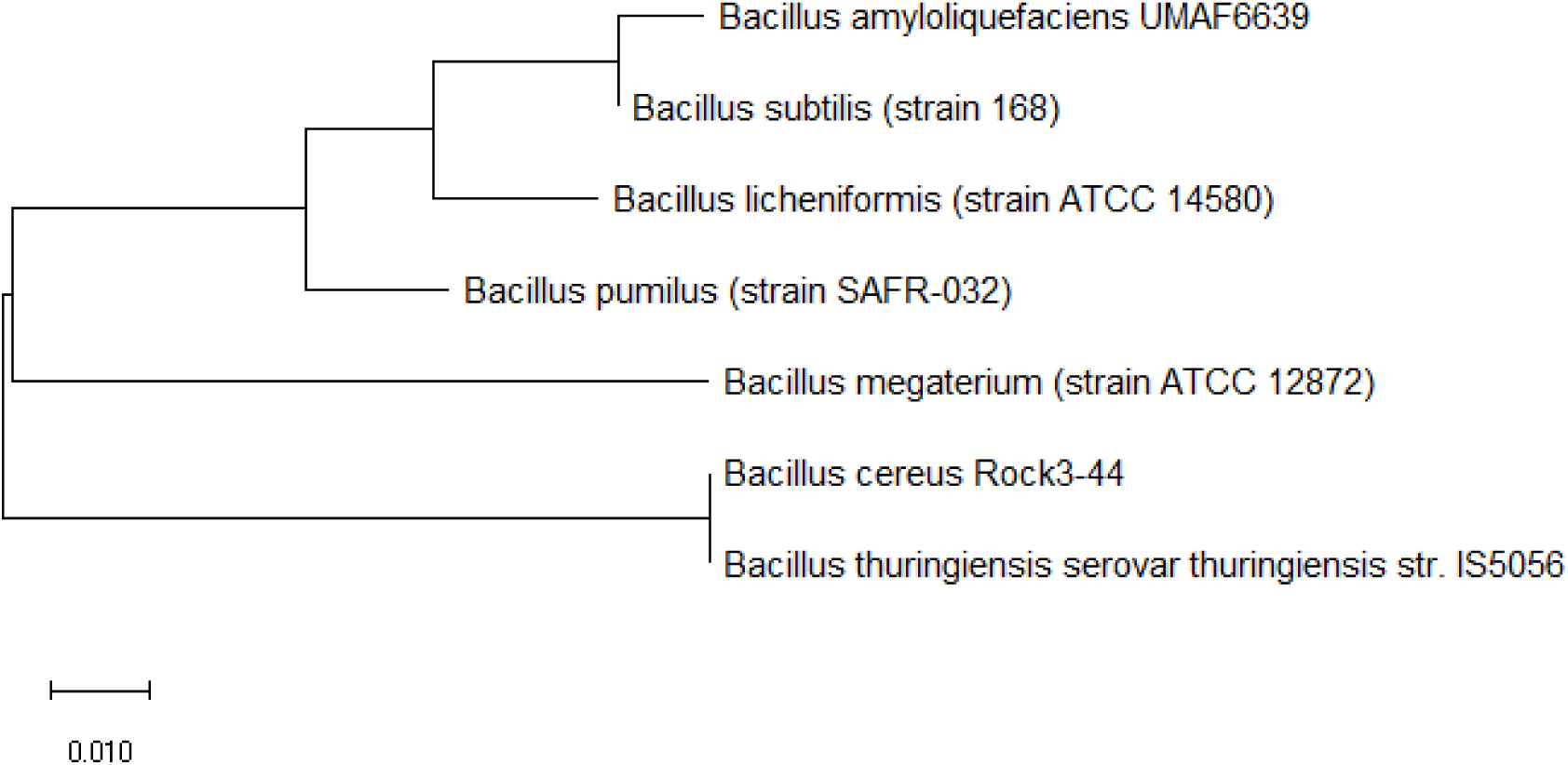
Maximum likelihood phylogenetic tree based on ribosomal protein L19 of different *Bacillus* species.

Figure 8 shows the maximum likelihood phylogenetic tree based on ribosomal protein L24 of different *Bacillus* species. The data revealed that ribosomal protein L24 could reproduce major branches of the 16S rRNA phylogenetic tree, and thus, it holds phylogenetic significance for explaining the evolutionary relatedness of different *Bacillus* species.

**Figure 8:**
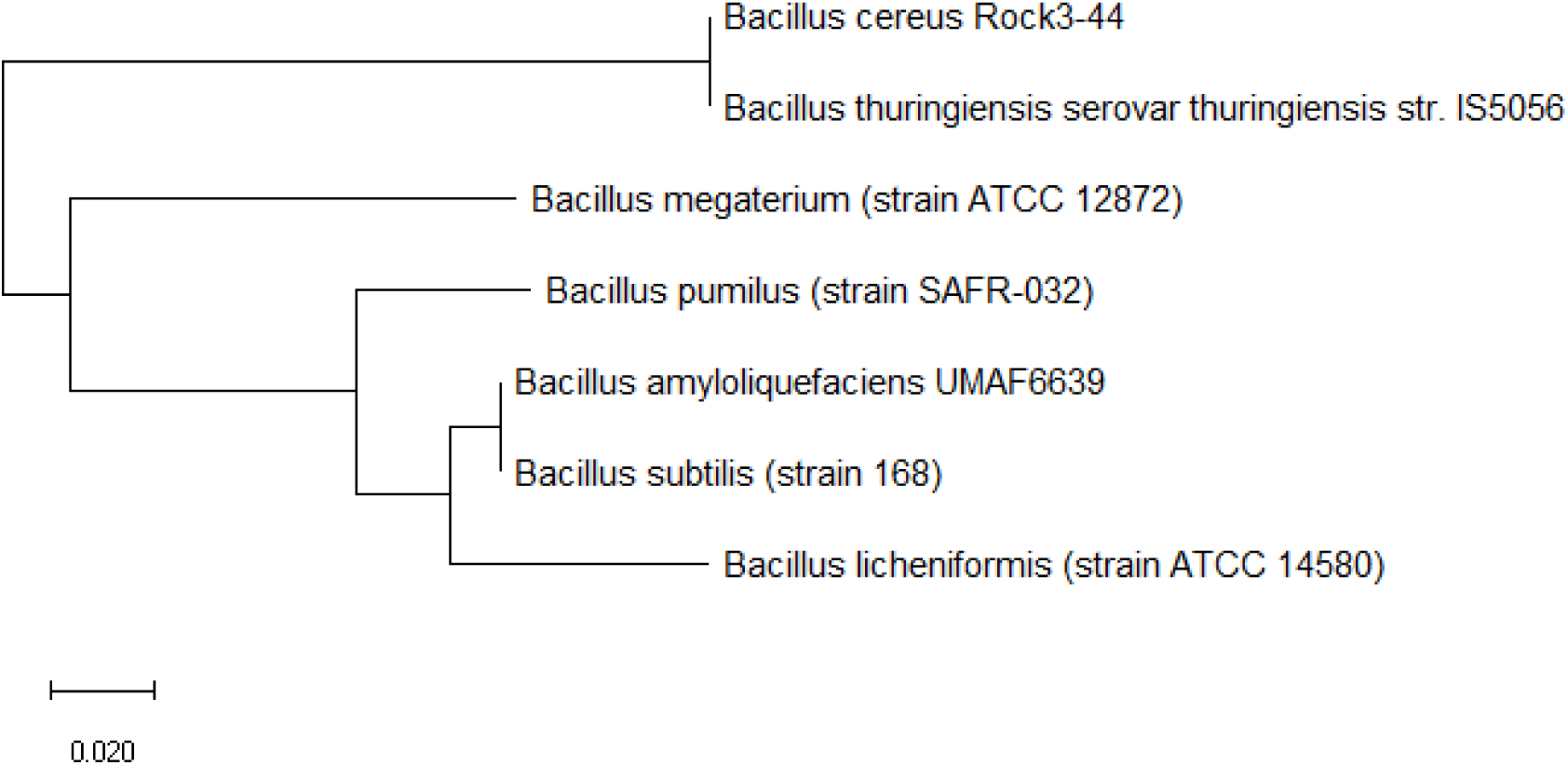
Maximum likelihood phylogenetic tree based on ribosomal protein L24 of different *Bacillus* species.

Figure 9 shows the maximum likelihood phylogenetic tree based on ribosomal protein L32 of different *Bacillus* species. The data revealed that ribosomal protein L32 could recapitulate the major branches of the 16S rRNA phylogenetic tree, and thus, the ribosomal protein holds phylogenetic significance in describing the evolutionary relationships between different *Bacillus* species.

**Figure 9:**
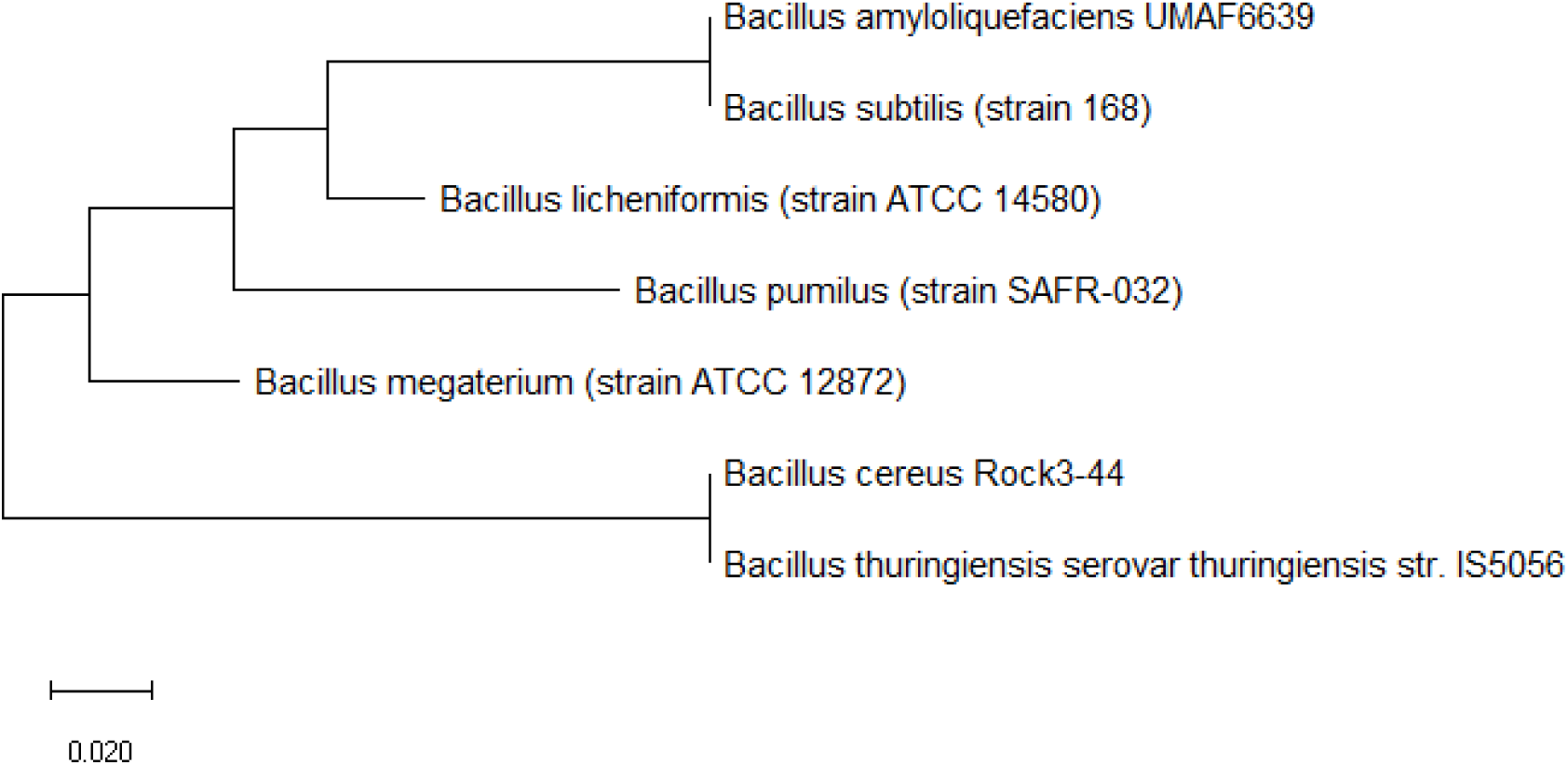
Maximum likelihood phylogenetic tree based on ribosomal protein L32 of different *Bacillus* species.

Figure 10 shows the maximum likelihood phylogenetic tree based on ribosomal protein S3 of different *Bacillus* species. The data revealed that ribosomal protein S3 could reproduce all the major branches of the 16S rRNA phylogenetic tree, and thus, it holds phylogenetic significance for explaining the evolutionary relationships between different *Bacillus* species.

**Figure 10:**
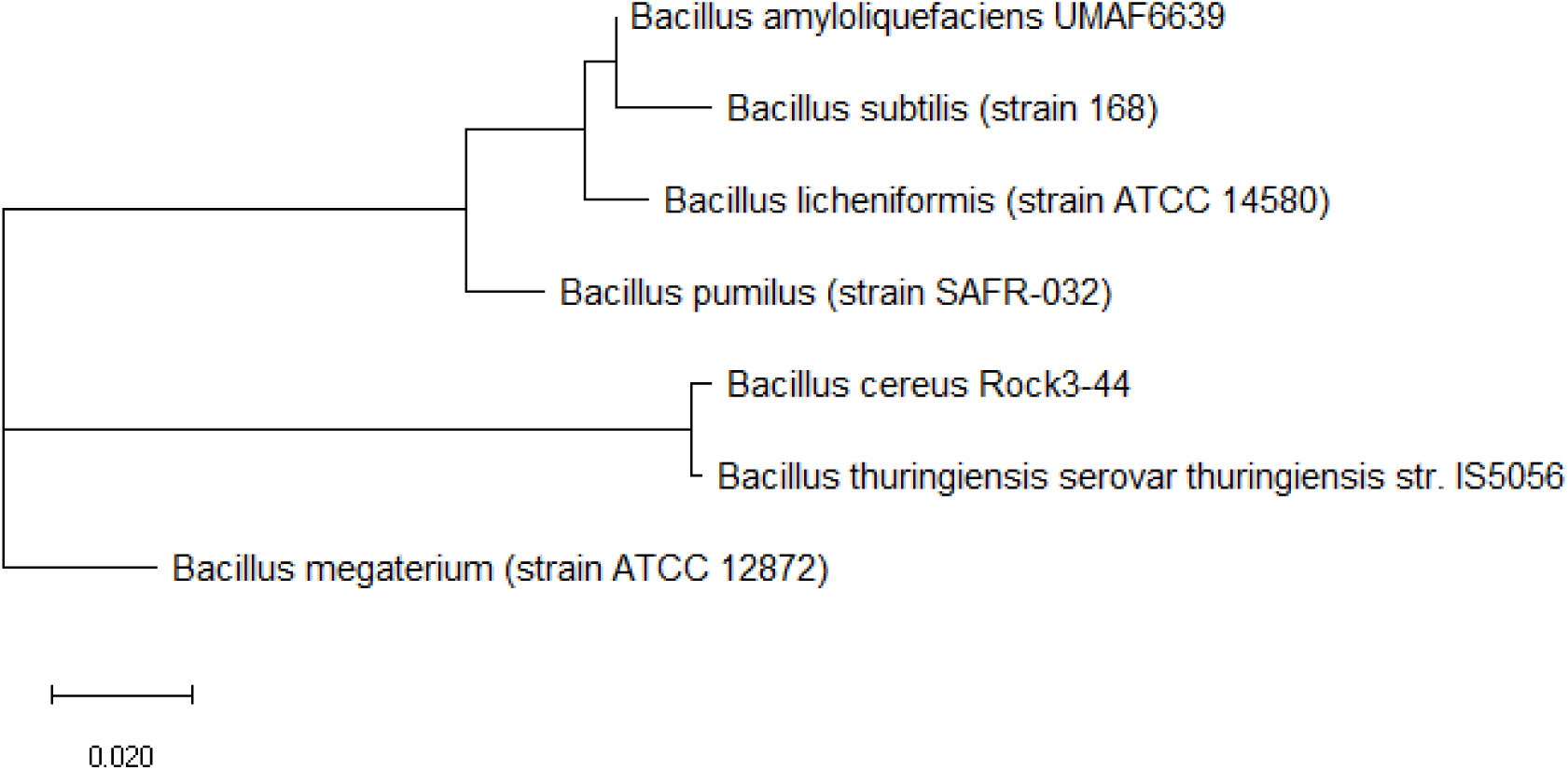
Maximum likelihood phylogenetic tree based on ribosomal protein S3 of different *Bacillus* species.

Figure 11 shows the maximum likelihood phylogenetic tree based on ribosomal protein S9 of different *Bacillus* species. The data revealed that ribosomal protein S9 could reproduce all the major branches of the 16S rRNA phylogenetic tree; thus, ribosomal protein S9 holds phylogenetic significance in explaining the evolutionary relatedness between different *Bacillus* species.

**Figure 11:**
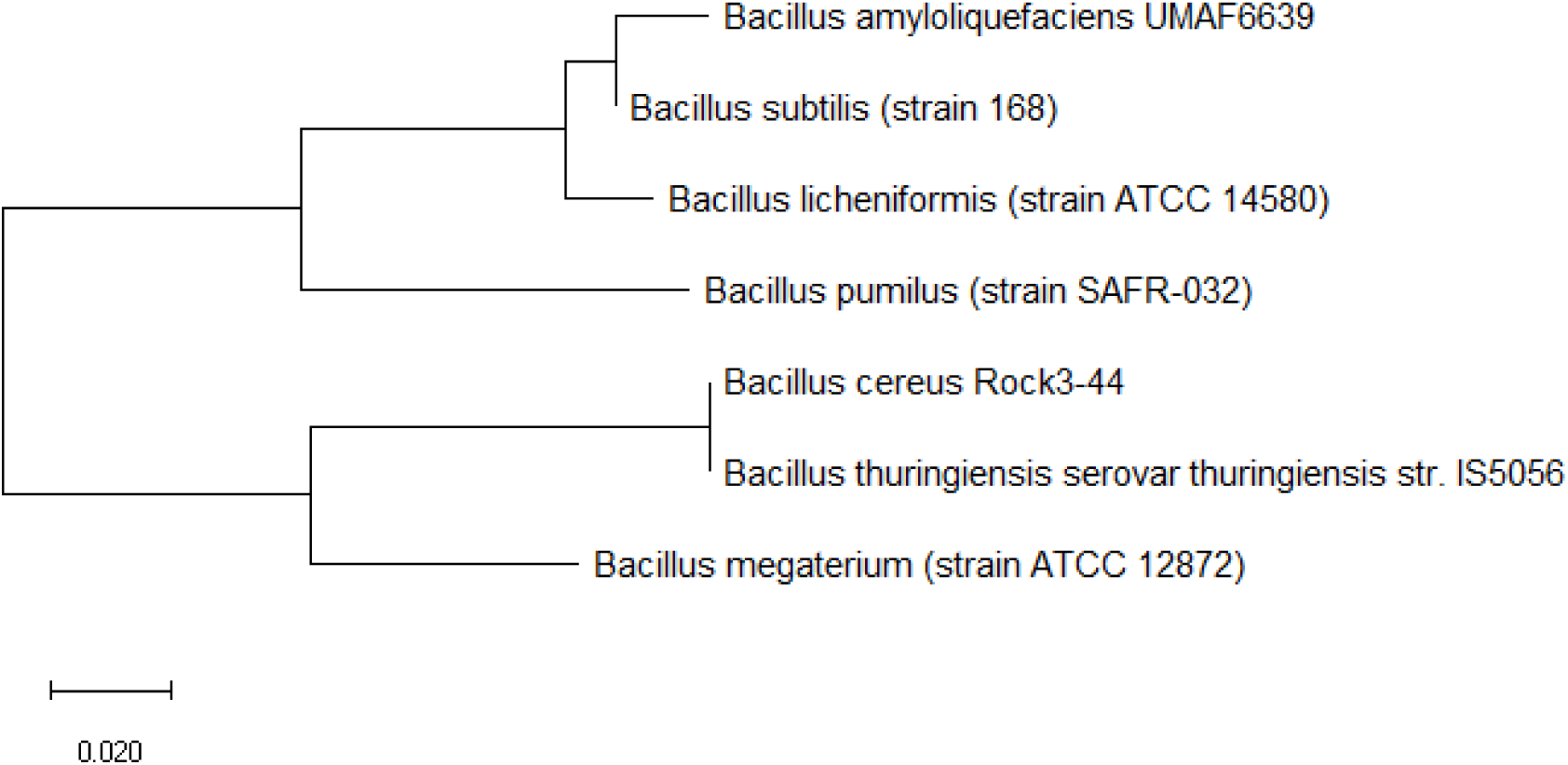
Maximum likelihood phylogenetic tree based on ribosomal protein S9 of different *Bacillus* species.

Figure 12 shows the maximum likelihood phylogenetic tree based on ribosomal protein S12 of different *Bacillus* species. The data revealed that ribosomal protein S12 could reproduce the major branches of the 16S rRNA phylogenetic tree of the different *Bacillus* species. Thus, ribosomal protein S12 holds phylogenetic significance for explaining the evolutionary relationships between different *Bacillus* species.

**Figure 12:**
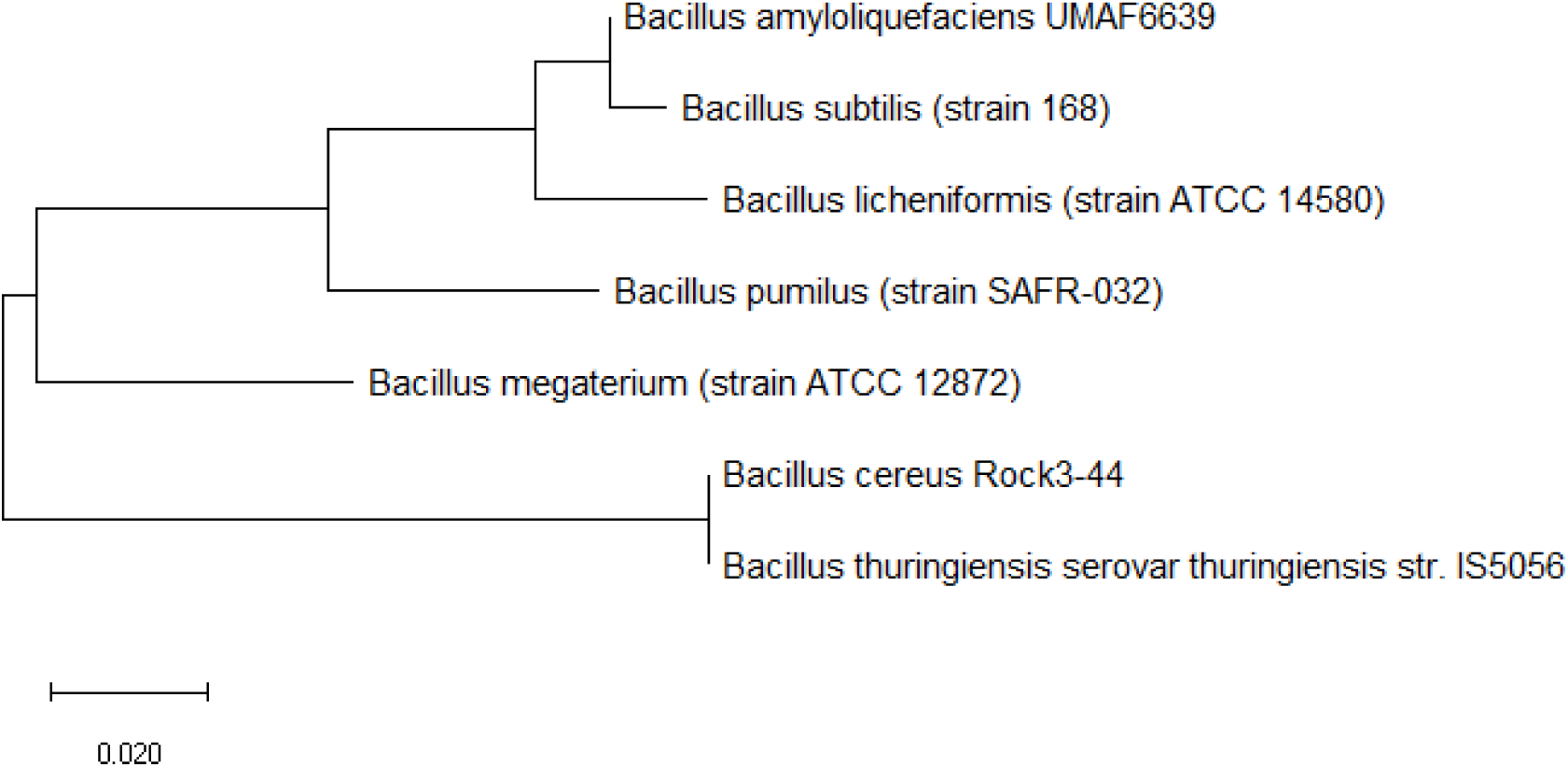
Maximum likelihood phylogenetic tree based on ribosomal protein S12 of different *Bacillus* species.

Figure 13 shows the maximum likelihood phylogenetic tree based on ribosomal protein S15 of different *Bacillus* species. The data revealed that ribosomal protein S15 could reproduce the major branches of the 16S rRNA phylogenetic tree for the *Bacillus* species investigated. Thus, ribosomal protein S15 holds phylogenetic significance for explaining the evolutionary relationships between different *Bacillus* species.

**Figure 13:**
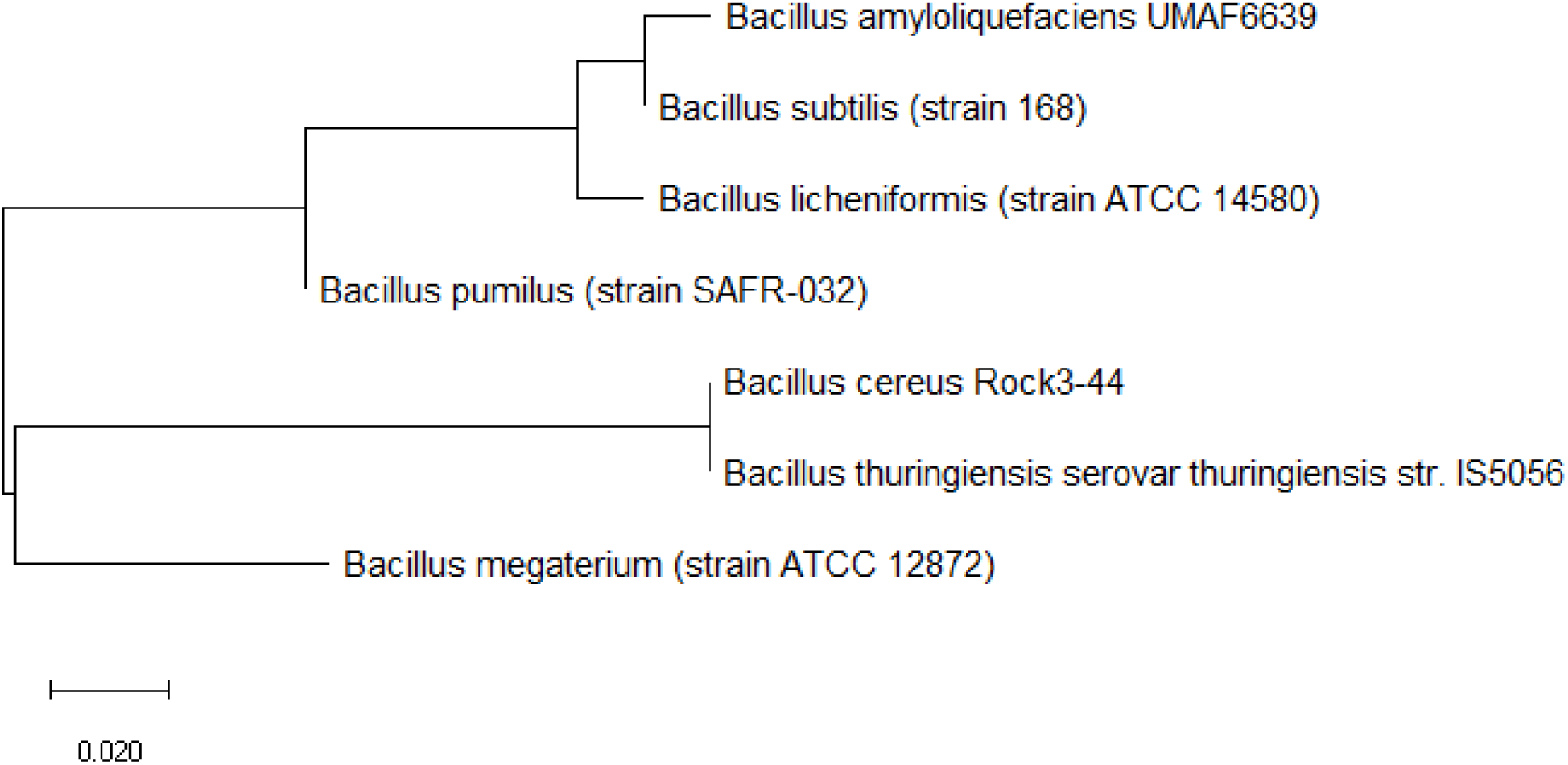
Maximum likelihood phylogenetic tree based on ribosomal protein S15 of different *Bacillus* species.

Figure 14 shows the maximum likelihood phylogenetic tree based on ribosomal protein S16 of different *Bacillus* species. The data revealed that the major branches of the 16S rRNA phylogenetic tree could be reproduced by ribosomal protein S16. Thus, ribosomal protein S16 holds phylogenetic significance in describing the phylogeny between the different *Bacillus* species.

**Figure 14:**
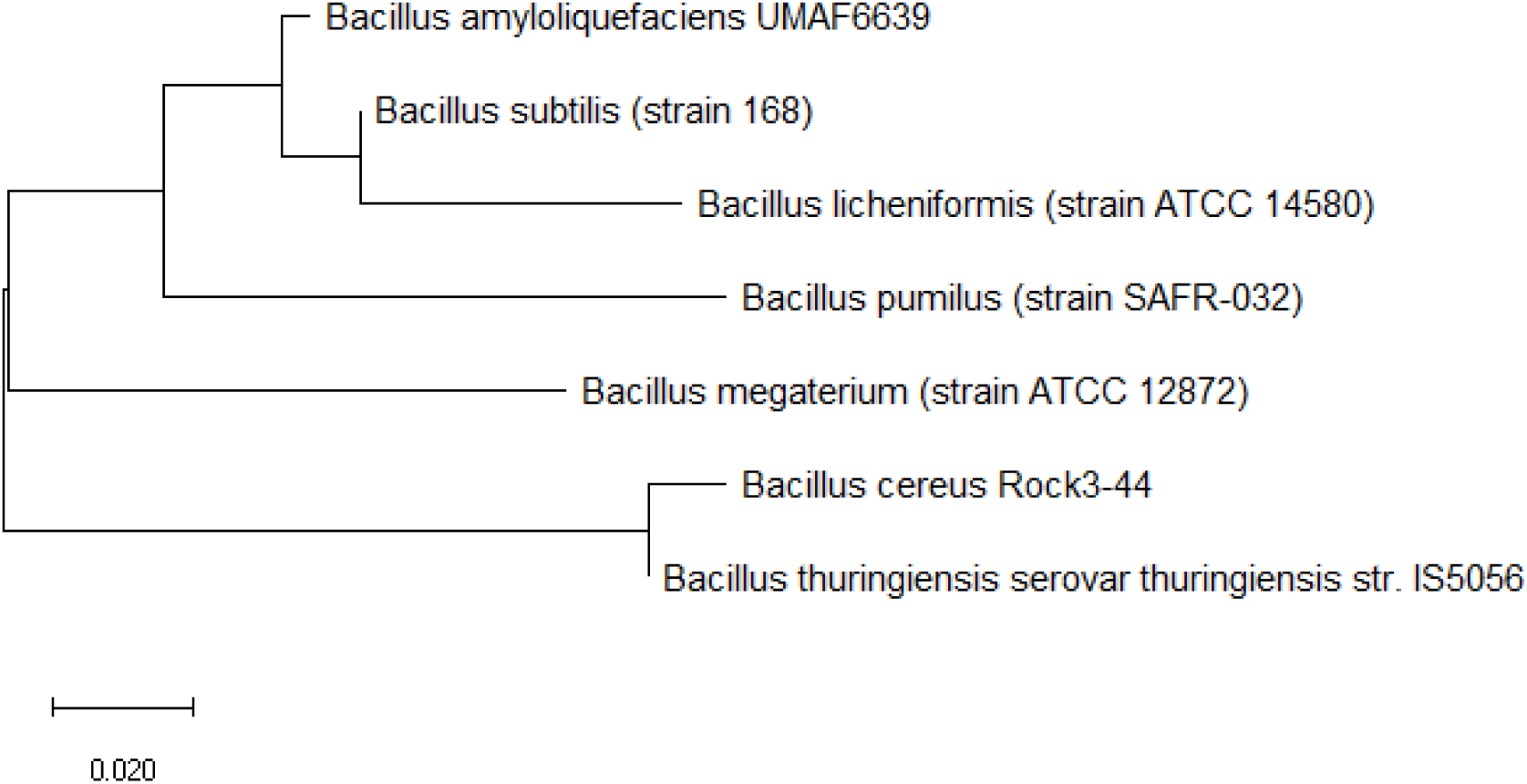
Maximum likelihood phylogenetic tree based on ribosomal protein S16 of different *Bacillus* species.

Figure 15 shows the maximum likelihood phylogenetic tree based on ribosomal protein S17 of different *Bacillus* species. The data revealed that ribosomal protein S17 could reproduce the major branches of the phylogenetic tree based on 16S rRNA. Thus, ribosomal protein S17 holds phylogenetic significance in describing the evolutionary relationship between different *Bacillus* species.

**Figure 15:**
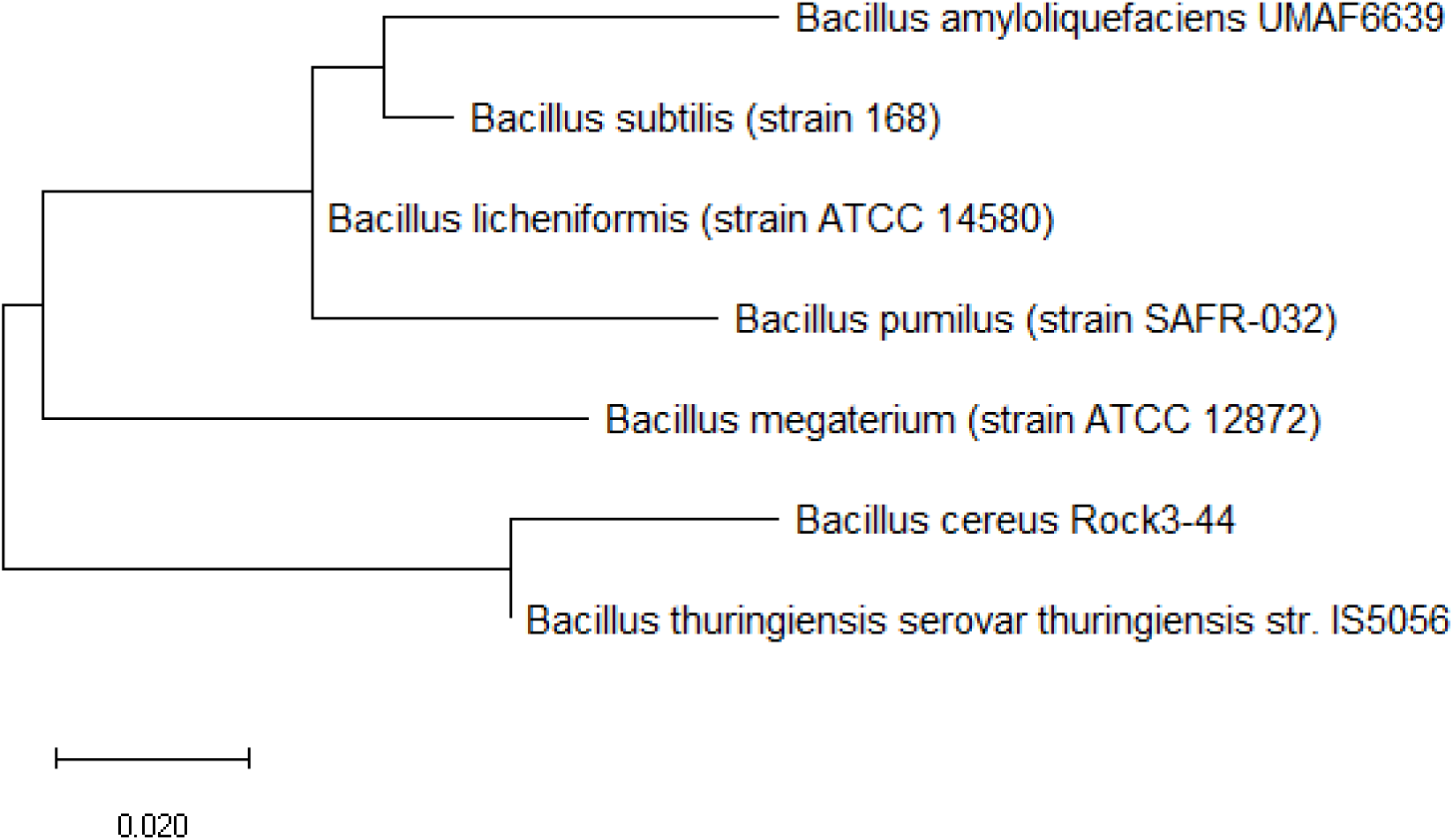
Maximum likelihood phylogenetic tree based on ribosomal protein S17 of different *Bacillus* species.

Figure 16 shows the maximum likelihood phylogenetic tree based on ribosomal protein S18 of different *Bacillus* species. The data revealed that ribosomal protein S18 could reproduce the major branches of the phylogenetic tree based on 16S rRNA. Thus, ribosomal protein S18 holds phylogenetic significance in explaining the phylogeny of different *Bacillus* species.

**Figure 16:**
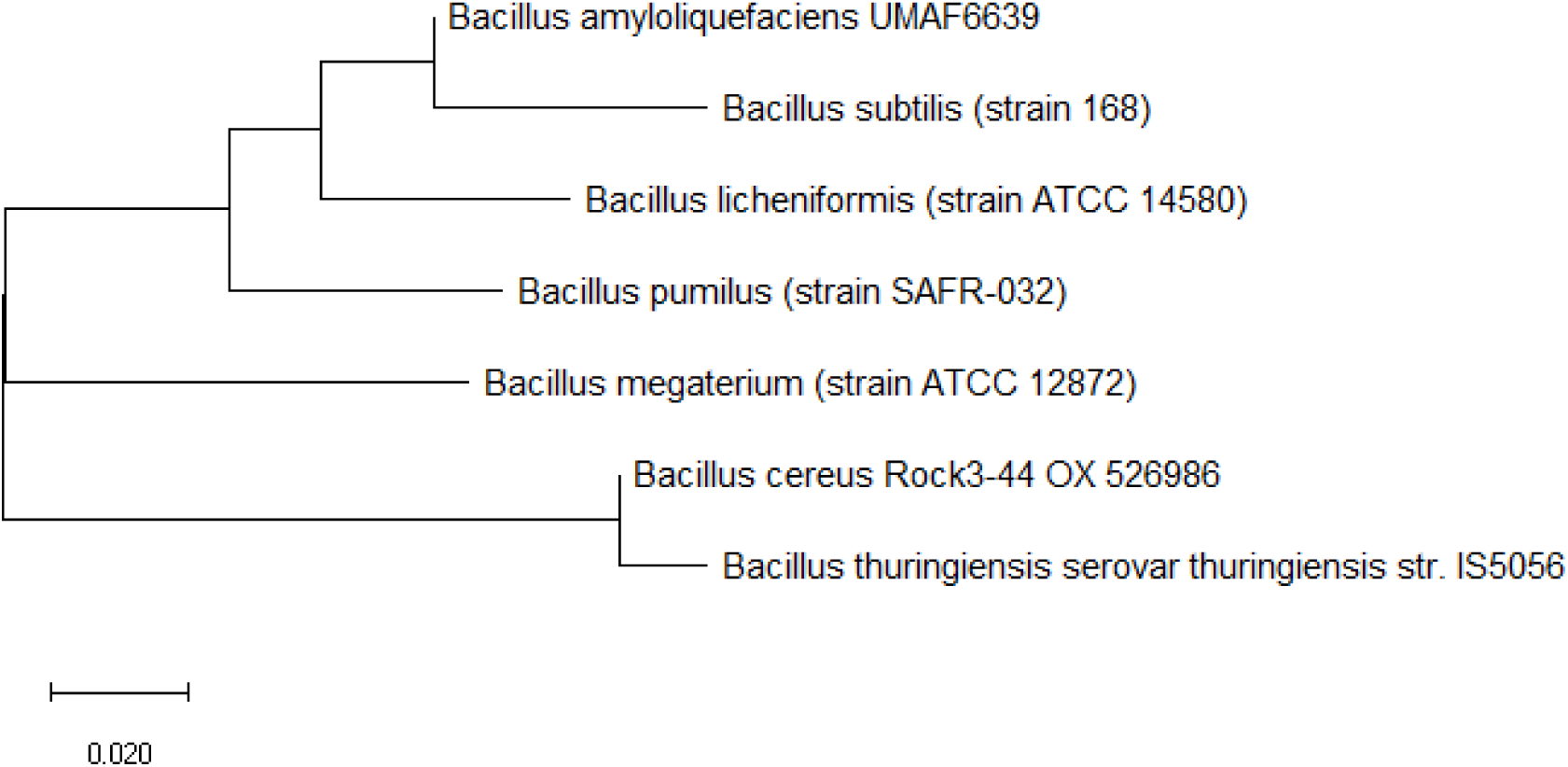
Maximum likelihood phylogenetic tree based on ribosomal protein S18 of different *Bacillus* species.

Overall, phylogenetic analysis revealed that many ribosomal proteins could explain the evolutionary relationships between the investigated *Bacillus* species. Specifically, data from Table 1 revealed that ribosomal protein L6, L7/12, L9, L13, L24, L32, S3, S9, S12, S15, S16, S17, and S18 could explain the phylogeny between the different *Bacillus* species well. Importantly, both ribosomal proteins from the large and small ribosome subunits were found to hold phylogenetic significance; thereby, indicating that both ribosome subunits and their proteins likely coevolved with each other. Coevolution of the ribosomal proteins that hold phylogenetic significance is likely given the evolutionary selection force that exerted pressure for simultaneous coevolution of the ribosomal proteins. Existence of coevolution between the ribosomal proteins would help explain why many ribosomal proteins in the large and small ribosome subunit could describe the evolutionary relationships between the different *Bacillus* species.

**Table 1:** Different categories of ribosomal proteins in *Bacillus* species

| Category | Ribosomal proteins |
| --- | --- |
| Hold phylogenetic significance | L6, L7/L12, L9, L13, L24, L32, S3, S9, S12, S15, S16, S17, S18 |
| Hold partial phylogenetic significance | L1, L2, L3, L4, L10, L11, L14, L15, L16, L18, L19, L21, L22, L29, L30, L31 Type B, L33, L35, S1, S4, S5, S6, S7, S8, S11, S13, S19, S20 |
| Too highly conserved | L5, L36, S2, S10, S21 |
| No phylogenetic significance | L7Ae, L17, L20, L23, L27, L28, L31, L34, S14 Type Z |

In addition to ribosomal proteins that hold complete phylogenetic significance in enabling the reproduction of the phylogenetic tree of the different *Bacillus* species described by 16S rRNA, there was a larger subset of ribosomal proteins that hold partial phylogenetic significance. These ribosomal proteins are typified by the general ability to reproduce major branches of the maximum likelihood phylogenetic tree that described the evolutionary relationships between different *Bacillus* species, but with the attendant inability to correctly place the phylogenetic position of 1 or 2 species. Specifically, the most commonly misplaced species were *Bacillus licheniformis* and *Bacillus pumilus*, which suggested that the phylogeny between the two species might be hard to disentangle using chronicled sequence information from ribosomal proteins. This group of ribosomal protein include ribosomal protein L22, L29, L30, L31 Type B, L33, L35, S1, S4, S5, S6, S7, S8, S11, S13, S19, S20.

Another class of ribosomal proteins were ones where the sequence was too highly conserved to allow them to encode meaningful evolutionary information in its sequence space. These ribosomal proteins came from both the small and large ribosome subunits and were ribosomal protein L5, L36, S2, S10, and S21. Lacking sequence diversity, maximum likelihood phylogenetic tree constructed based on these ribosomal proteins were unable to disentangle the evolutionary history of the different *Bacillus* species.

Ribosomal proteins unable to inform the phylogenetic relationships between the different *Bacillus* species form the final class of ribosomal proteins. Similarly, these ribosomal proteins came from both the small and large ribosome subunits and were ribosomal protein L7Ae, L17, L20, L23, L27, L28, L31, L34 and S14 Type Z. Maximum likelihood phylogenetic tree constructed using amino acid sequence information from these ribosomal proteins, in general, could not reproduce the phylogenetic positions of many species as compared to the phylogenetic tree based on 16S rRNA.

## Conclusions

As part of the ribosome that participates in protein translation, ribosomal proteins are highly conserved in sequence and structure given the strict evolutionary pressure exerted on them for retaining the essential folds and structural motifs critical for enzymatic and structural functions. However, success with using 16S rRNA as a biomarker for understanding the evolutionary trajectories of different species across the three domains of life revealed that ribosomal proteins and ribosomal RNA could be endowed with sufficient sequence diversity to help chronicle the evolutionary history of respective species. Thus, individual ribosomal proteins could be useful in understanding the phylogeny of different species within and between genera if they are endowed with sufficient sequence diversity to chronicle the impacts of different selection pressure on the species throughout evolutionary timescales.

By determining whether individual ribosomal protein could reproduce the phylogenetic tree described by 16S rRNA, this study sought to assess if individual ribosomal proteins could chronicle the evolutionary history of the investigated *Bacillus* species. Results indicated that depending on whether the ribosomal proteins could reproduce the 16S rRNA phylogenetic tree of the *Bacillus* species, four different groups of ribosomal proteins could be delineated. The first group represents those that hold phylogenetic significance in reproducing the phylogenetic positions of the *Bacillus* species according to the phylogeny depicted by 16S rRNA. This group of ribosomal proteins include ribosomal protein L6, L7/12, L9, L13, L24, L32, S3, S9, S12, S15, S16, S17, and S18. In addition to those that could reproduce accurately all the phylogenetic positions of the *Bacillus* species, there was another group of ribosomal proteins that hold partial phylogenetic significance. Specifically, they could reproduce major branches of the 16S rRNA phylogenetic tree but differ in the placement of one or two *Bacillus* species. This group of ribosomal proteins include ribosomal protein L22, L29, L30, L31 Type B, L33, L35, S1, S4, S5, S6, S7, S8, S11, S13, S19, S20.

Ribosomal proteins whose amino acid sequence was too highly conserved to allow those to encode the evolutionary history of the different *Bacillus* species also existed. These ribosomal proteins lump the different *Bacillus* species into one or two phylogenetic group. Ribosomal proteins in this group include ribosomal protein L5, L36, S2, S10, and S21. Finally, there were also ribosomal proteins that did not hold any phylogenetic significance where they randomly placed the different *Bacillus* species along the branches of the reconstructed phylogenetic tree. This group of ribosomal proteins comprises ribosomal protein L7Ae, L17, L20, L23, L27, L28, L31, L34 and S14 Type Z.

Overall, depending on their sequence diversity, ribosomal proteins exhibit varying utility in reproducing the phylogeny of different *Bacillus* species. However, many ribosomal proteins could reproduce the 16S rRNA phylogenetic tree of the *Bacillus* species under study and revealed that evolutionary processes must be at work influencing the evolutionary trajectories of some of the ribosomal proteins nestled within the large and small subunit of the ribosome. Interestingly, ribosomal proteins that hold phylogenetic significance came from both the large and small subunit of the ribosome, which implied that protein-protein as well as protein-rRNA interactions could be at work influencing the evolutionary trajectories of the ribosome structure and function along species line. Specifically, the points of contact between proteins and between proteins and rRNA could serve as points at which selection pressure are exerted on the ribosomal proteins. Collectively, specific ribosomal proteins could inform the phylogeny of different *Bacillus* species.

## Supplementary materials

Additional data on ribosomal proteins with partial or no phylogenetic significance in differentiating members of the *Bacillus* genus probed in this study is encapsulated as a supplementary file

## Conflicts of interest

The author declares no conflicts of interest.

## Funding

No funding was used in this work.

## Notes

### Competing Interest Statement

The authors have declared no competing interest.

## References

1 Yang, B., Wang, Y. & Qian, P.-Y. Sensitivity and correlation of hypervariable regions in 16S rRNA genes in phylogenetic analysis. BMC Bioinformatics 17, 135, doi: 10.1186/s12859-016-0992-y (2016).

2 Janda, J. M. & Abbott, S. L. 16S rRNA Gene Sequencing for Bacterial Identification in the Diagnostic Laboratory: Pluses, Perils, and Pitfalls. Journal of Clinical Microbiology 45, 2761, doi: 10.1128/JCM.01228-07 (2007).

3 Ng, W. Annotation of ribosomal protein mass peaks in MALDI-TOF mass spectra of bacterial species and their phylogenetic significance. bioRxiv, 2020.2007.2015.203653, doi: 10.1101/2020.07.15.203653 (2020).

4 Nakamura, S. et al. Ribosomal subunit protein typing using matrix-assisted laser desorption ionization time-of-flight mass spectrometry (MALDI-TOF MS) for the identification and discrimination of Aspergillus species. BMC Microbiology 17, 100, doi: 10.1186/s12866-017-1009-3 (2017).

5 Singhal, N., Kumar, M., Kanaujia, P. K. & Virdi, J. S. MALDI-TOF mass spectrometry: an emerging technology for microbial identification and diagnosis. Frontiers in Microbiology 6, 791 (2015).

6 Li, Y. et al. Application of MALDI-TOF MS to rapid identification of anaerobic bacteria. BMC Infectious Diseases 19, 941, doi: 10.1186/s12879-019-4584-0 (2019).

7 Fernández-Álvarez, C., Torres-Corral, Y. & Santos, Y. Use of ribosomal proteins as biomarkers for identification of Flavobacterium psychrophilum by MALDI-TOF mass spectrometry. Journal of Proteomics 170, 59–69, doi: https://doi.org/10.1016/j.jprot.2017.09.007 (2018).

8 Suarez, S. et al. Ribosomal proteins as biomarkers for bacterial identification by mass spectrometry in the clinical microbiology laboratory. Journal of Microbiological Methods 94, 390–396, doi: https://doi.org/10.1016/j.mimet.2013.07.021 (2013).

9 Tomachewski, D. et al. Ribopeaks: a web tool for bacterial classification through m/z data from ribosomal proteins. Bioinformatics 34, 3058–3060, doi: 10.1093/bioinformatics/bty215 (2018).

10 Roberts, E., Sethi, A., Montoya, J., Woese, C. R. & Luthey-Schulten, Z. Molecular signatures of ribosomal evolution. Proceedings of the National Academy of Sciences 105, 13953–13958, doi: 10.1073/pnas.0804861105 (2008).

11 Shu, L.-J. & Yang, Y.-L. Bacillus Classification Based on Matrix-Assisted Laser Desorption Ionization Time-of-Flight Mass Spectrometry—Effects of Culture Conditions. Scientific Reports 7, 15546, doi: 10.1038/s41598-017-15808-5 (2017).

12 Quast, C. et al. The SILVA ribosomal RNA gene database project: improved data processing and web-based tools. Nucleic Acids Research 41, D590–D596, doi: 10.1093/nar/gks1219 (2012).

13 Hall, B. G. Building Phylogenetic Trees from Molecular Data with MEGA. Molecular Biology and Evolution 30, 1229–1235, doi: 10.1093/molbev/mst012 (2013).

