## Supplementary materials for "Ribosomal proteins could explain the phylogeny of *Bacillus* species"

Wenfa Ng

Department of Chemical and Biomolecular Engineering, National University of Singapore,

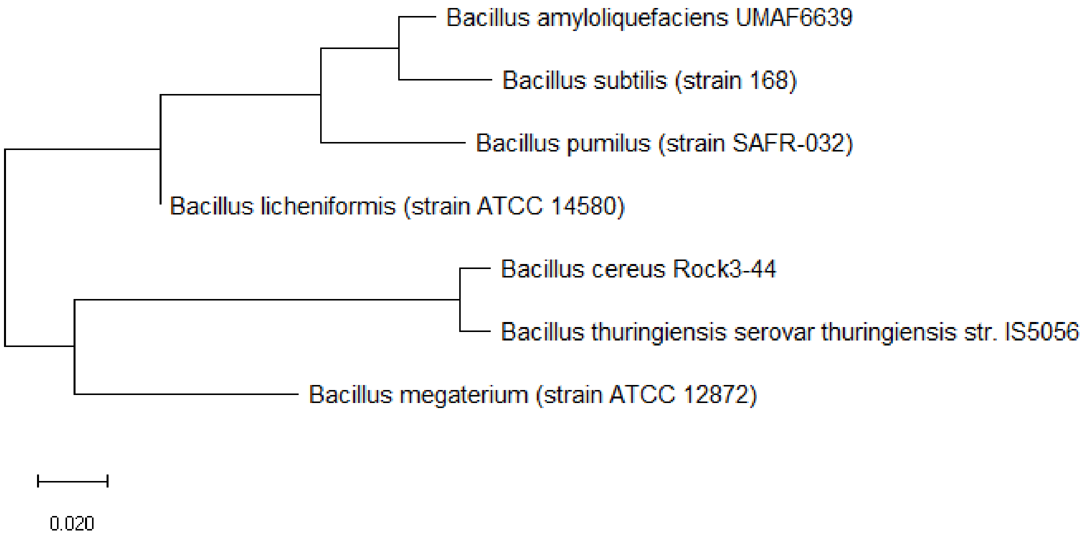


**Figure S1:** Maximum likelihood phylogenetic tree based on ribosomal protein L1 of different *Bacillus* species.

Figure S1 shows the maximum likelihood phylogenetic tree based on ribosomal protein L1 of different *Bacillus* species. Observations of the phylogenetic tree revealed that ribosomal protein L1 could explain the phylogeny of the investigated *Bacillus* species well except for the misplacement of the phylogenetic relationship between *B. licheniformis* and *B. pumilus*. Thus, ribosomal protein L1 holds partial phylogenetic significance.

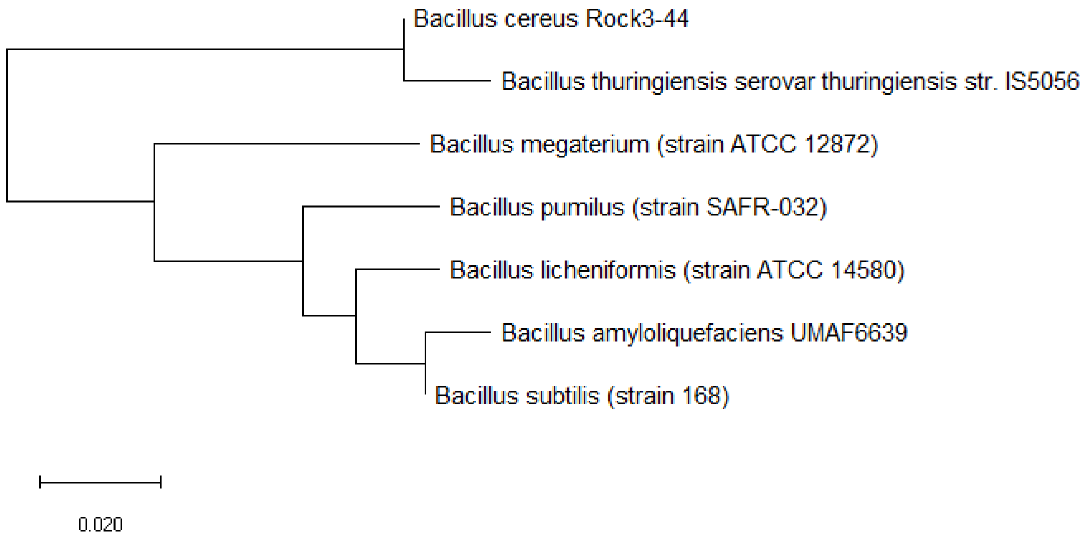


**Figure S2:** Maximum likelihood phylogenetic tree based on ribosomal protein L2 of different *Bacillus* species.

The maximum likelihood phylogenetic tree based on ribosomal protein L2 of different *Bacillus* species is shown in Figure S2. Specifically, the phylogenetic tree revealed the misplacement of the position of *B. megaterium*, while other branches of the phylogenetic tree showed high concordance with that based on 16S rRNA. Thus, ribosomal protein L2 holds partial phylogenetic significance in explaining the phylogeny of *Bacillus* species.

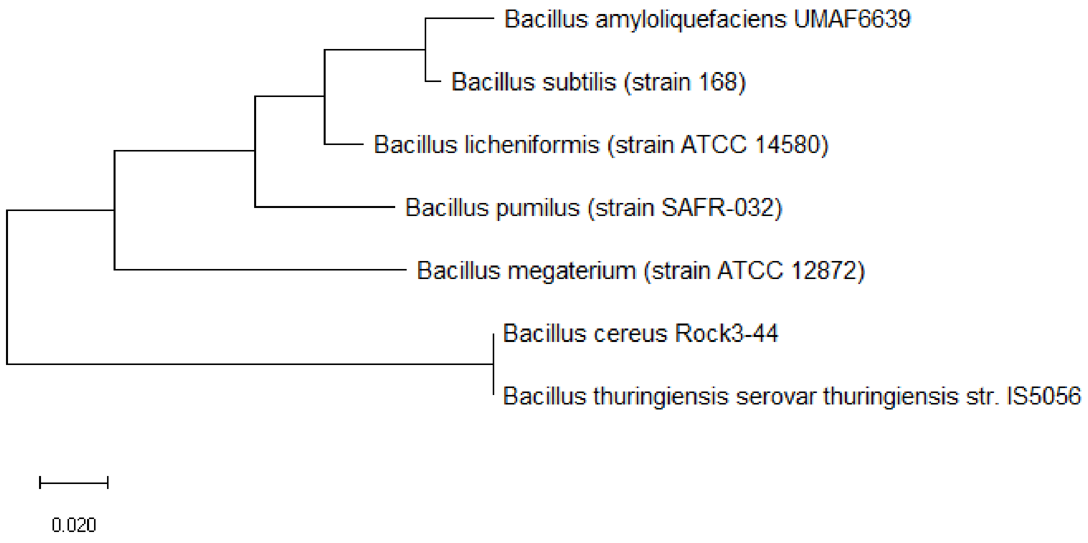

**Figure S3:** Maximum likelihood phylogenetic tree based on ribosomal protein L3 of different *Bacillus* species.

Figure S3 shows the maximum likelihood phylogenetic tree based on ribosomal protein L3 of different *Bacillus* species. Similar to the case for ribosomal protein L2, the phylogenetic tree based on ribosomal protein L3 misplaced the position of *B. megaterium* while other branches of the phylogenetic tree showed high concordance with that of 16S rRNA. Hence, ribosomal protein L3 holds partial phylogenetic significance in explaining the phylogeny of *Bacillus* species.

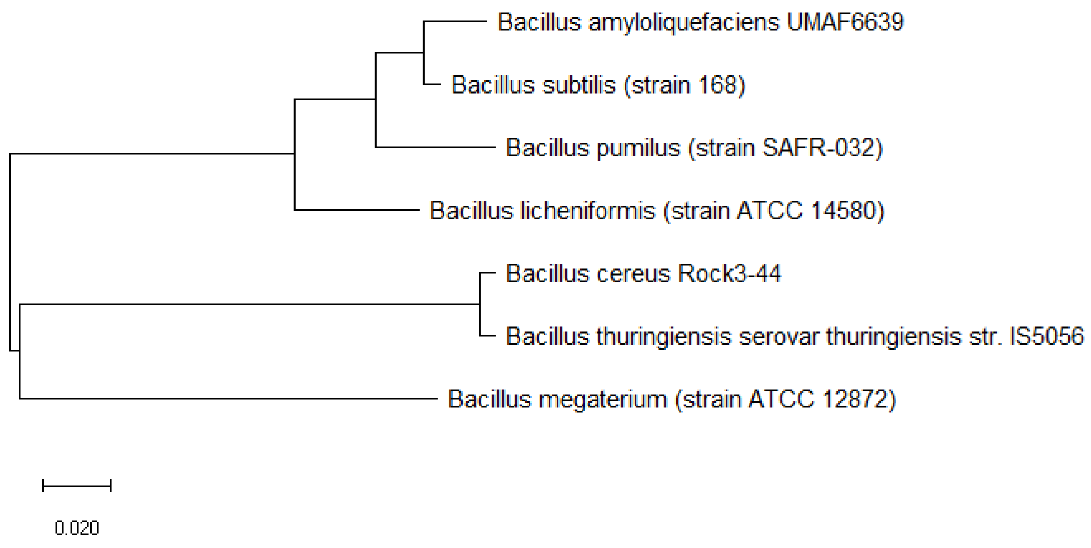


**Figure S4:** Maximum likelihood phylogenetic tree based on ribosomal protein L4 of different *Bacillus* species

Figure S4 shows the maximum likelihood phylogenetic tree based on ribosomal protein L4 of different *Bacillus* species. The data revealed that ribosomal protein L4 could not explain the phylogenetic relationship between *B. licheniformis* and *B. pumilus* well, resulting in the misplacement of their phylogenetic positions in the tree. Thus, ribosomal protein L4 could only hold partial phylogenetic significance in explaining the phylogeny of *Bacillus* species.

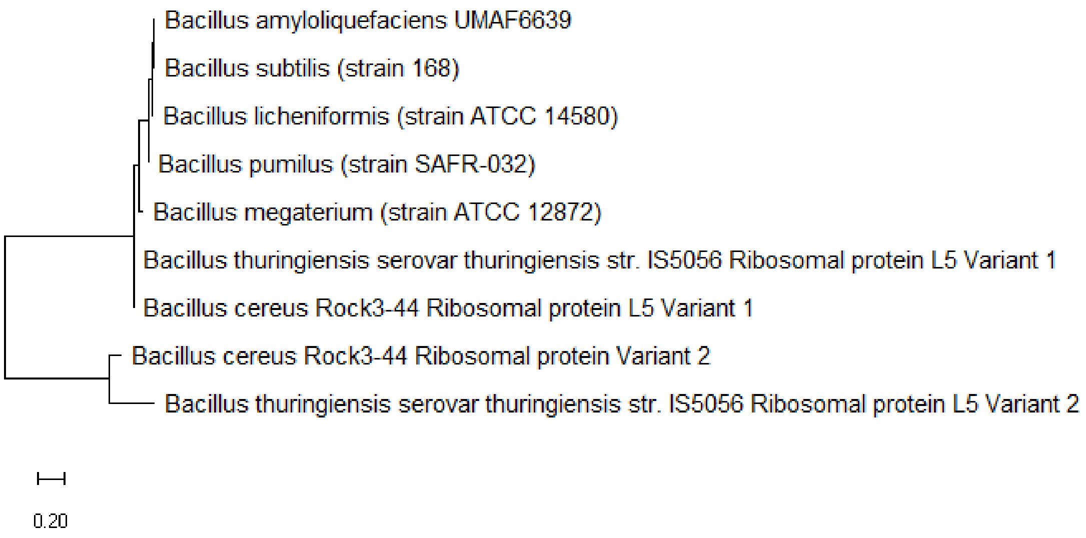


**Figure S5:** Maximum likelihood phylogenetic tree based on ribosomal protein L5 of different *Bacillus* species.

Figure S5 shows the maximum likelihood phylogenetic tree based on ribosomal protein L5 of different *Bacillus* species. The data revealed that ribosomal protein L5 of different *Bacillus* species were highly conserved in sequence and could not chronicle the evolutionary history of the different *Bacillus* species. Thus, ribosomal protein L5 does not hold phylogenetic significance for the investigated *Bacillus* species.

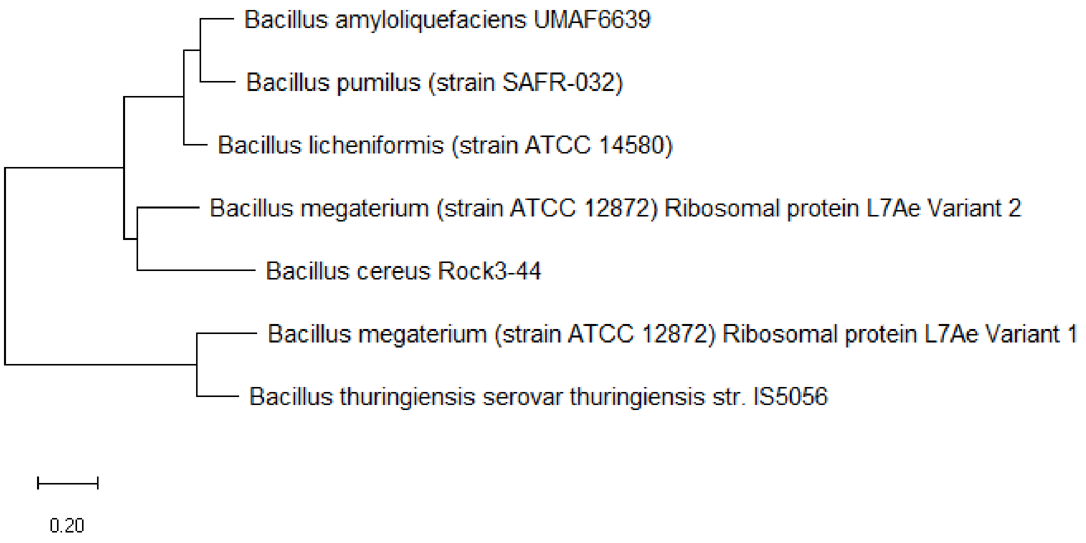


**Figure S6:** Maximum likelihood phylogenetic tree based on ribosomal protein L7Ae of different *Bacillus* species.

Figure S6 shows the maximum likelihood phylogenetic tree based on ribosomal protein L7Ae of different *Bacillus* species. Specifically, it could be seen that ribosomal protein L7Ae could not reproduce major branches of the 16S rRNA phylogenetic tree for the same *Bacillus* species. Thus, ribosomal protein L7Ae does not hold phylogenetic significance.

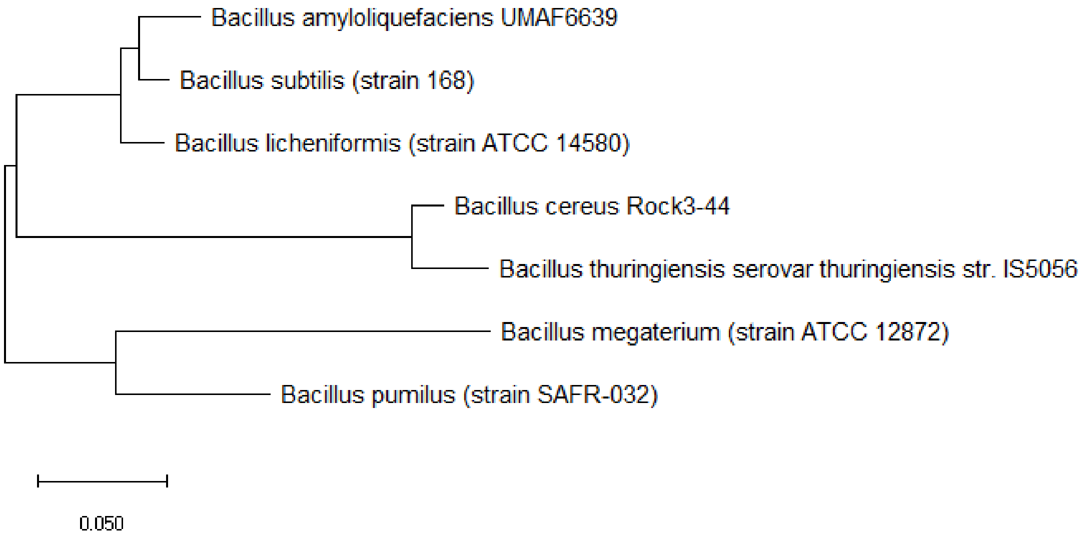


**Figure S7:** Maximum likelihood phylogenetic tree based on ribosomal protein L10 of different *Bacillus* species.

Figure S7 shows the maximum likelihood phylogenetic tree based on ribosomal protein L10 of different *Bacillus* species. The data revealed that except for *Bacillus pumilus*, all major branches of the 16S rRNA phylogenetic tree for the same species could be reproduced by that based on ribosomal protein L10. Thus, ribosomal protein L10 hold partial phylogenetic significance for the set of *Bacillus* species investigated.

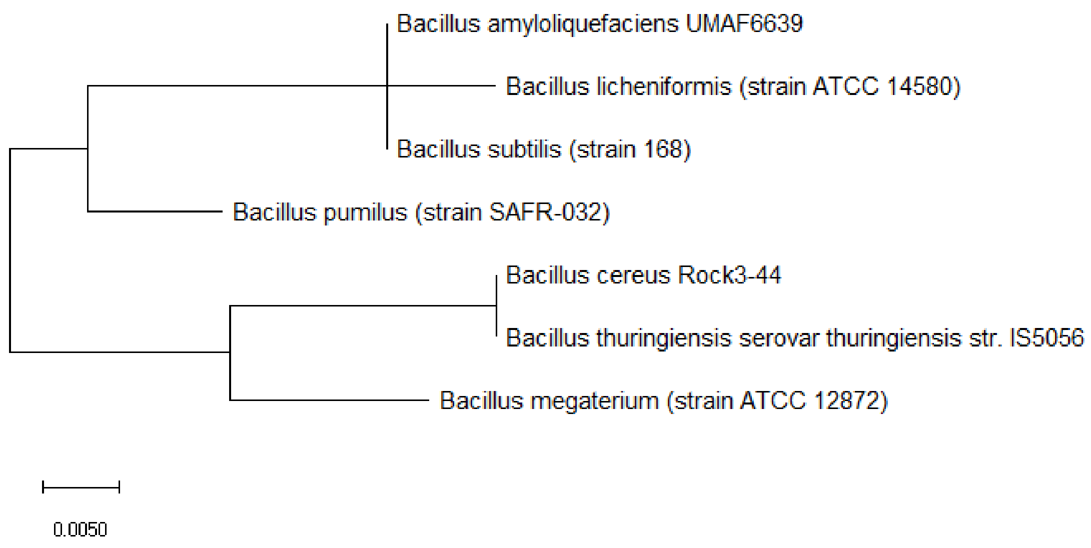


**Figure S8:** Maximum likelihood phylogenetic tree based on ribosomal protein L11 of different *Bacillus* species.

Figure S8 shows the maximum likelihood phylogenetic tree based on ribosomal protein L11 of different *Bacillus* species. The data revealed that the phylogenetic placement of *Bacillus licheniformis* was wrong when compared with that in the 16S rRNA phylogenetic tree. Thus, ribosomal protein L11 hold partial phylogenetic significance for explaining the evolutionary relatedness of different *Bacillus* species.

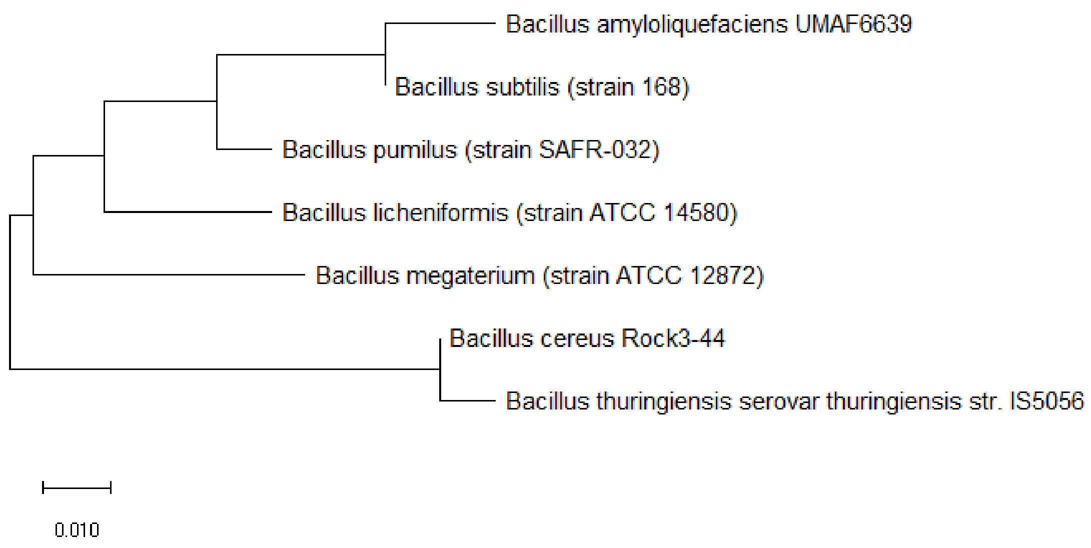

**Figure S9:** Maximum likelihood phylogenetic tree based on ribosomal protein L14 of different *Bacillus* species.

Figure S9 shows the maximum likelihood phylogenetic tree based on ribosomal protein L14 of different *Bacillus* species. The data revealed that except for the wrong placement of *Bacillus pumilus*, ribosomal protein L14 could reproduce the major branches of the 16S rRNA phylogenetic tree. Thus, ribosomal protein L14 hold partial phylogenetic significance for the investigated set of *Bacillus* species.

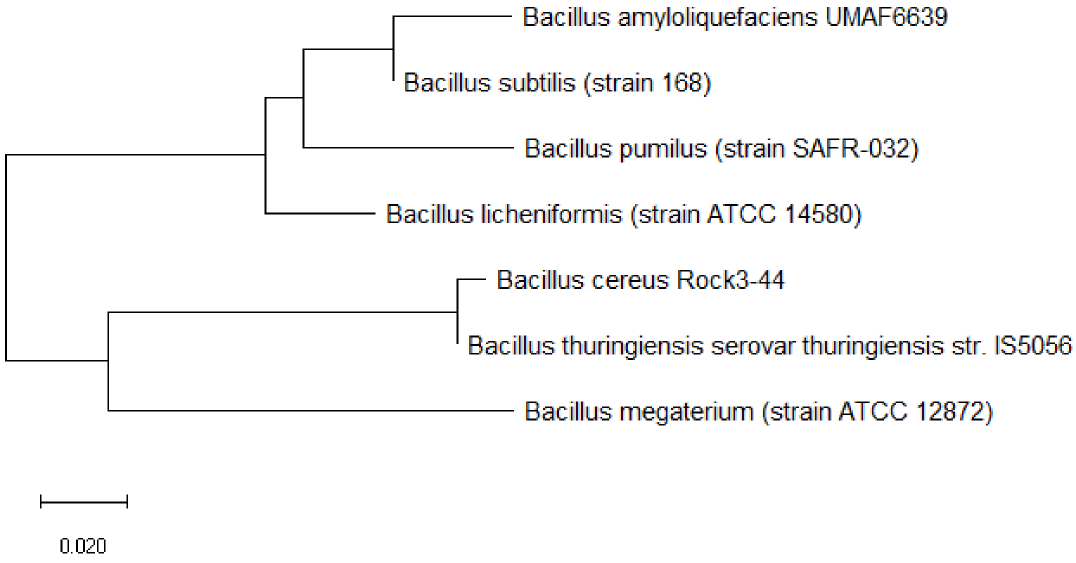


**Figure S10:** Maximum likelihood phylogenetic tree based on ribosomal protein L15 of different *Bacillus* species.

Figure S10 shows the maximum likelihood phylogenetic tree based on ribosomal protein L15 of different *Bacillus* species. Similar to the case for ribosomal protein L14, ribosomal protein L15 placed the phylogenetic position of *Bacillus pumilus* wrongly, but could, in general, reproduce major branches of the 16S rRNA phylogenetic tree. Thus, ribosomal protein L14 holds partial phylogenetic significance in explaining the phylogeny of the different *Bacillus* species.

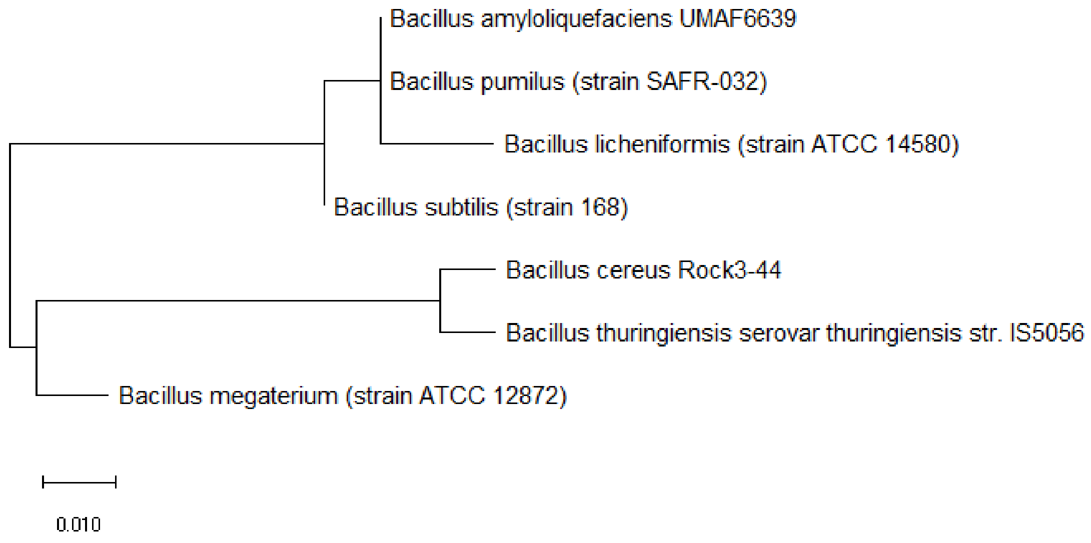


**Figure S11:** Maximum likelihood phylogenetic tree based on ribosomal protein L16 of different *Bacillus* species.

Figure S11 shows the maximum likelihood phylogenetic tree based on ribosomal protein L16 of different *Bacillus* species. The data revealed that ribosomal protein L16 placed the phylogenetic positions of *Bacillus subtilis* and *Bacillus pumilus* wrongly. Thus, ribosomal protein L16 holds partial phylogenetic significance for explaining the phylogeny of the different *Bacillus* species.

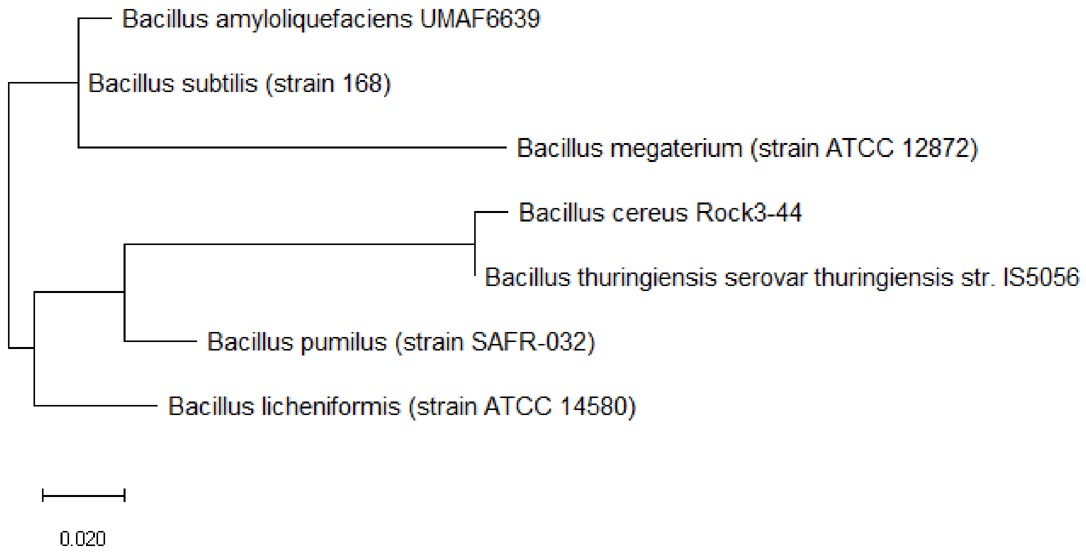

**Figure S12:** Maximum likelihood phylogenetic tree based on ribosomal protein L17 of different *Bacillus* species.

Figure S12 shows the maximum likelihood phylogenetic tree based on ribosomal protein L17 of different *Bacillus* species. The data revealed that ribosomal protein L17 could not place the phylogenetic positions of *Bacillus megaterium*, *Bacillus pumilus*, and *Bacillus licheniformis* accurately as compared to that based on 16S rRNA. Thus, ribosomal protein L17 does not hold phylogenetic significance for explaining the evolutionary relatedness of different *Bacillus* species.

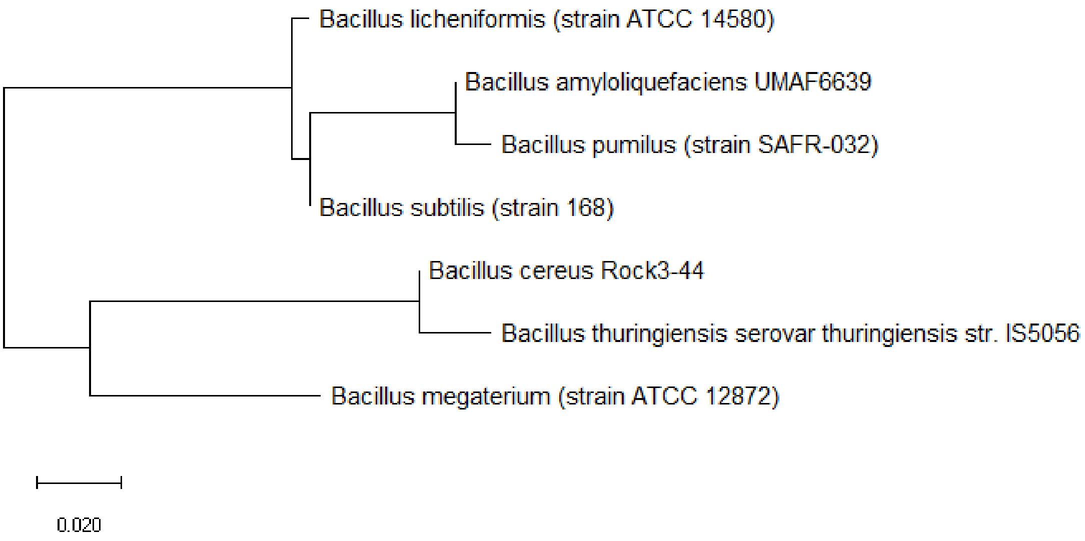


**Figure S13:** Maximum likelihood phylogenetic tree based on ribosomal protein L20 of different *Bacillus* species.

Figure S13 shows the maximum likelihood phylogenetic tree based on ribosomal protein L20 of different *Bacillus* species. The data revealed that ribosomal protein L20 could not place *Bacillus subtilis*, *Bacillus licheniformis*, and *Bacillus pumilus* at the correct positions of the 16S rRNA phylogenetic tree of the same *Bacillus* species. Thus, ribosomal protein L20 does not hold phylogenetic significance for the set of *Bacillus* species.

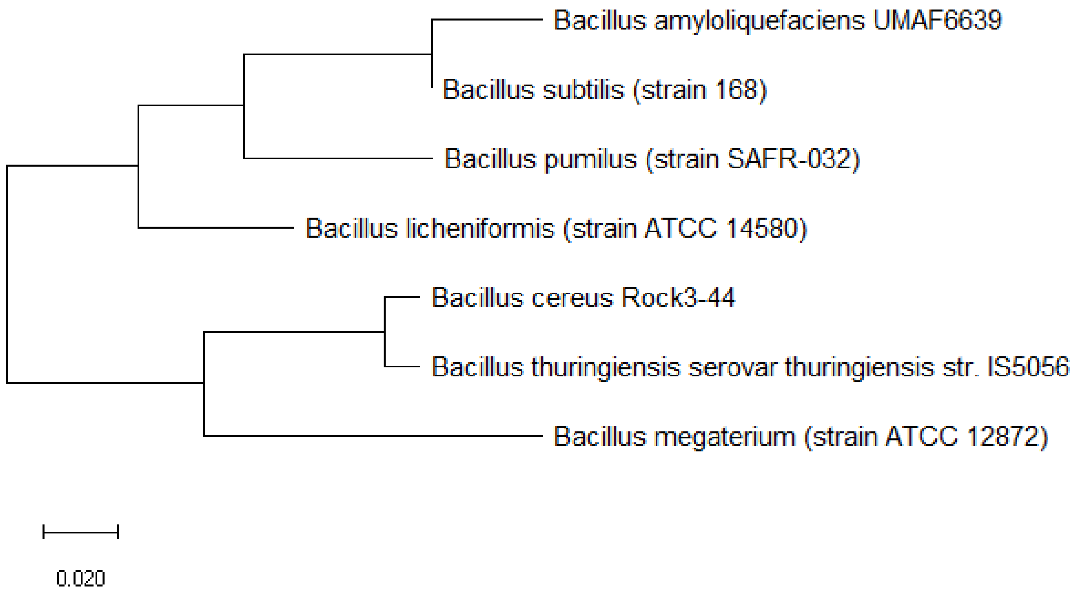


**Figure S14:** Maximum likelihood phylogenetic tree based on ribosomal protein L21 of different *Bacillus* species.

Figure S14 shows the maximum likelihood phylogenetic tree based on ribosomal protein L21 of different *Bacillus* species. Ribosomal protein L21 could reproduce the major branches of the 16S rRNA phylogenetic tree of the *Bacillus* species, but placed the relative positions of *Bacillus pumilus* and *Bacillus licheniformis* incorrectly. Thus, ribosomal protein L21 hold partial phylogenetic significance for this set of *Bacillus* species.

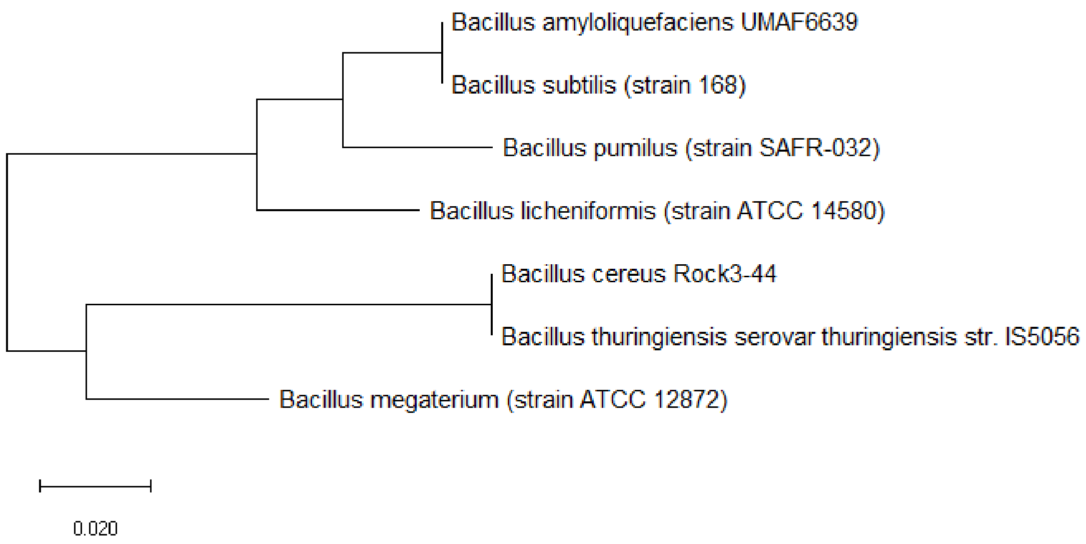

**Figure S15:** Maximum likelihood phylogenetic tree based on ribosomal protein L22 of different *Bacillus* species.

Figure S15 shows the maximum likelihood phylogenetic tree based on ribosomal protein L22 of different *Bacillus* species. The data revealed that major branches of the 16S rRNA phylogenetic tree for this set of *Bacillus* species could be reproduced, but *Bacillus pumilus* and *Bacillus licheniformis* were incorrectly positioned. Thus, ribosomal protein L22 hold partial phylogenetic significance in explaining the phylogeny of this set of *Bacillus* species.

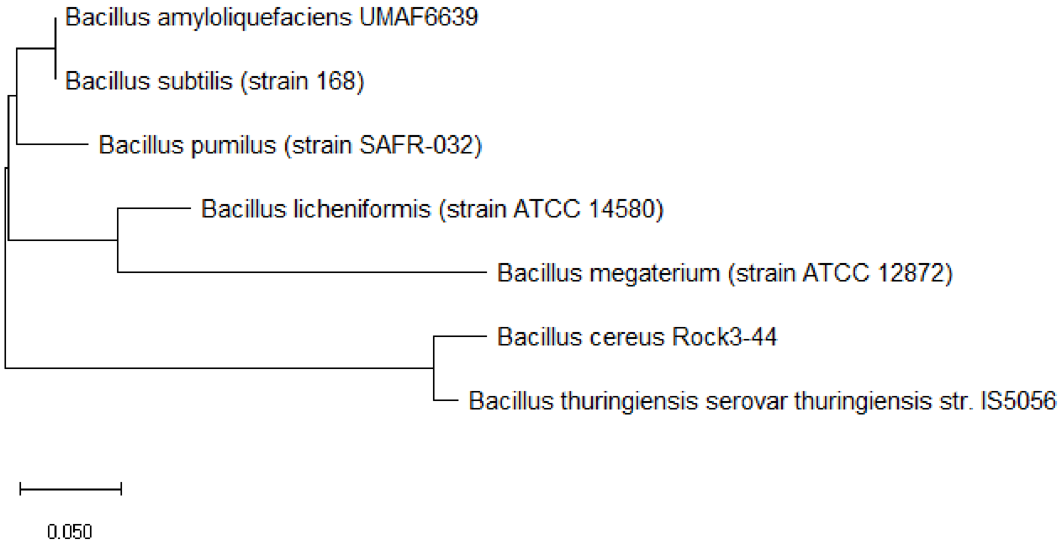


**Figure S16:** Maximum likelihood phylogenetic tree based on ribosomal protein L23 of different *Bacillus* species.

Figure S16 shows the maximum likelihood phylogenetic tree based on ribosomal protein L23 of different *Bacillus* species. The data revealed that ribosomal protein L23 could not place the phylogenetic positions of *Bacillus megaterium, Bacillus licheniformis*, and *Bacillus pumilus* correctly according to the phylogenetic tree based on 16S rRNA. Thus, ribosomal protein L23 does not hold phylogenetic significance for this set of *Bacillus* species.

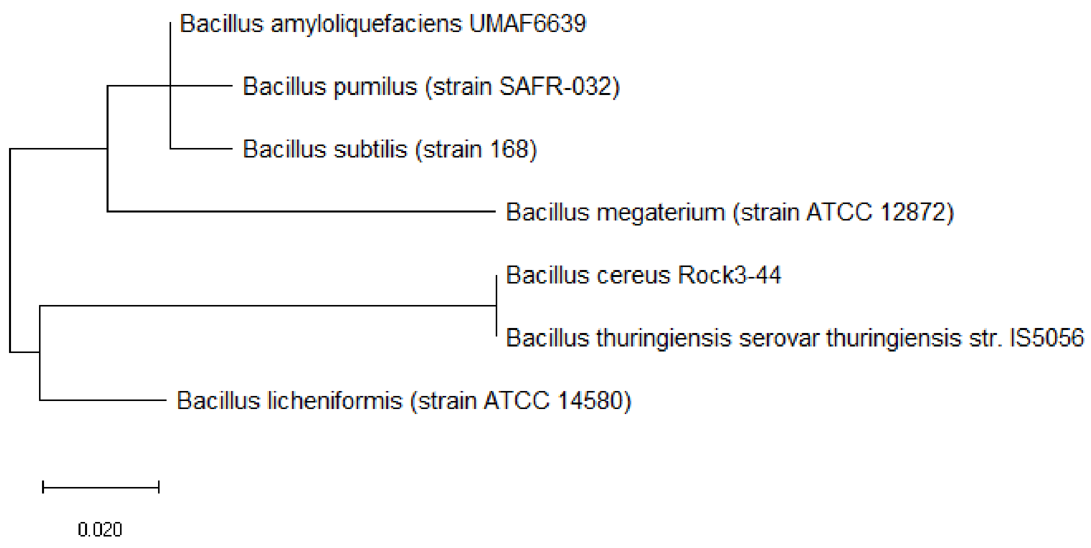


**Figure S17:** Maximum likelihood phylogenetic tree based on ribosomal protein L27 of different *Bacillus* species.

Figure S17 shows the maximum likelihood phylogenetic tree based on ribosomal protein L27 of different *Bacillus* species. The data revealed that ribosomal protein L27 could not place *Bacillus licheniformis*, *Bacillus megaterium*, and *Bacillus pumilus* at the correct phylogenetic positions with respect to the 16S rRNA phylogenetic tree. Thus, ribosomal protein L27 does not hold phylogenetic significance.

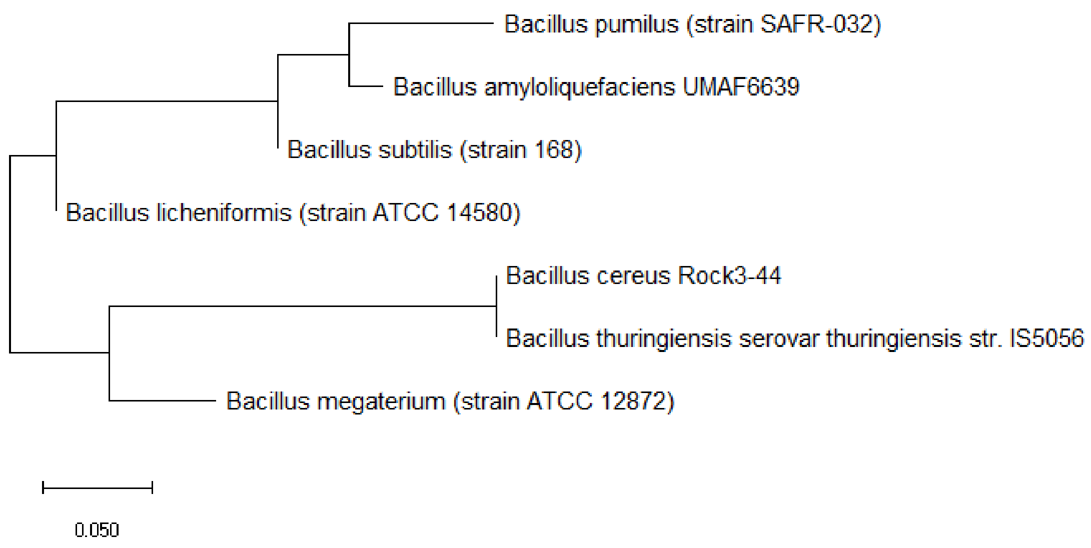

**Figure S18:** Maximum likelihood phylogenetic tree based on ribosomal protein L28 of different *Bacillus* species.

Figure S18 shows the maximum likelihood phylogenetic tree based on ribosomal protein L28 of different *Bacillus* species. The data revealed that ribosomal protein L28 could not place *Bacillus licheniformis* and *Bacillus pumilus* at the correct phylogenetic position with respect to the phylogeny described by 16S rRNA. Thus, ribosomal protein L28 does not hold phylogenetic significance in describing the evolutionary relationships of the different *Bacillus* species.

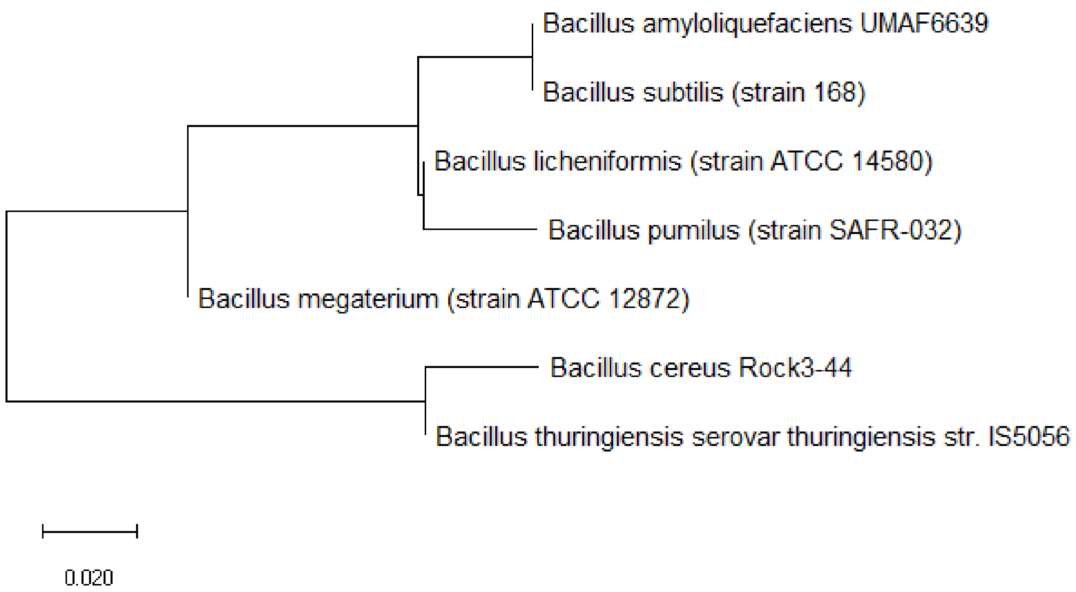


**Figure S19:** Maximum likelihood phylogenetic tree based on ribosomal protein L29 of different *Bacillus* species.

Figure S19 shows the maximum likelihood phylogenetic tree based on ribosomal protein L29 of different *Bacillus* species. The data revealed that ribosomal protein L29 could reproduce major branches of the 16S rRNA phylogenetic tree for the set of *Bacillus* species. However, the phylogenetic distances between different species were wrongly depicted. Thus, ribosomal protein L29 only hold partial phylogenetic significance for explaining the evolutionary relationships between the different *Bacillus* species.

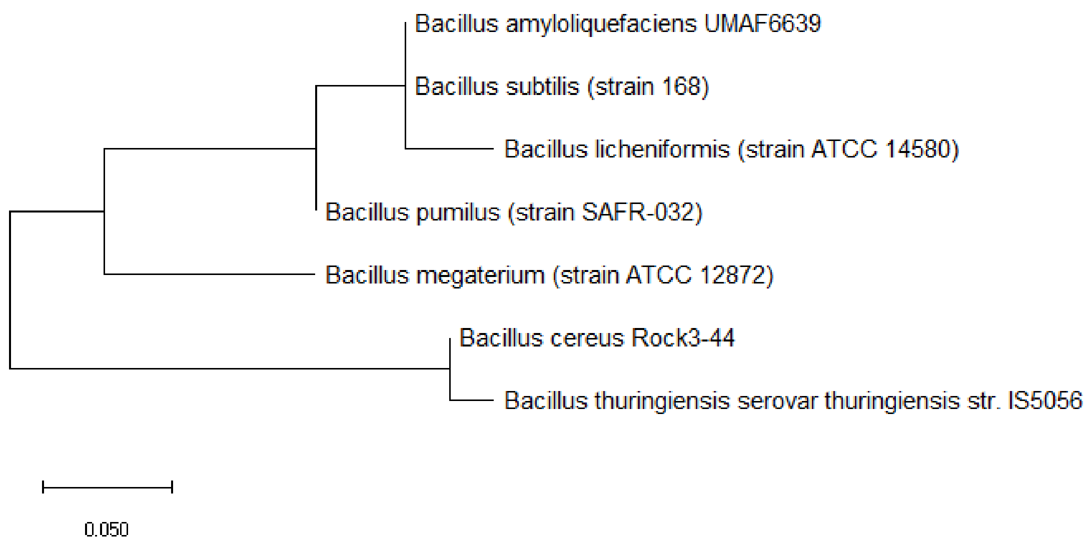


**Figure S20:** Maximum likelihood phylogenetic tree based on ribosomal protein L30 of different *Bacillus* species.

Figure S20 shows the maximum likelihood phylogenetic tree based on ribosomal protein L30 of different *Bacillus* species. The data revealed that ribosomal protein L30 could reproduce the major branches of the 16S rRNA phylogenetic tree but the phylogenetic distances between some species were wrongly depicted. Thus, ribosomal protein L30 only holds partial phylogenetic significance for the set of *Bacillus* species.

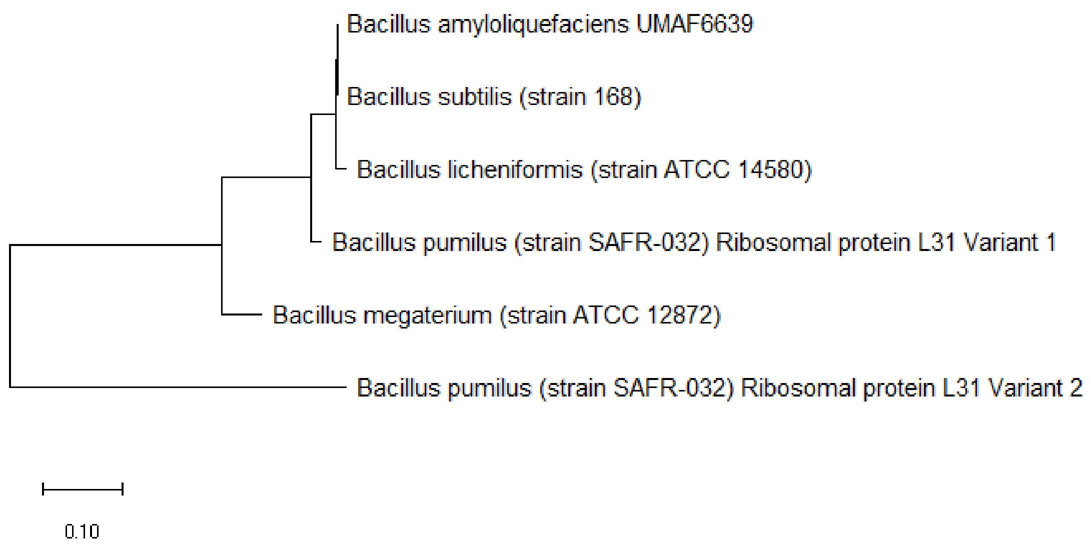

**Figure S21:** Maximum likelihood phylogenetic tree based on ribosomal protein L31 of different *Bacillus* species.

Figure S21 shows the maximum likelihood phylogenetic tree based on ribosomal protein L31 of different *Bacillus* species. The data revealed that ribosomal protein L31 could not differentiate the phylogenetic relationships between *Bacillus subtilis*, *Bacillus licheniformis*, and *Bacillus amyloliquefaciens* due to high conservation in amino acid sequence between different species. Thus, ribosomal protein L31 does not hold phylogenetic significance for explaining the phylogeny between different *Bacillus* species.

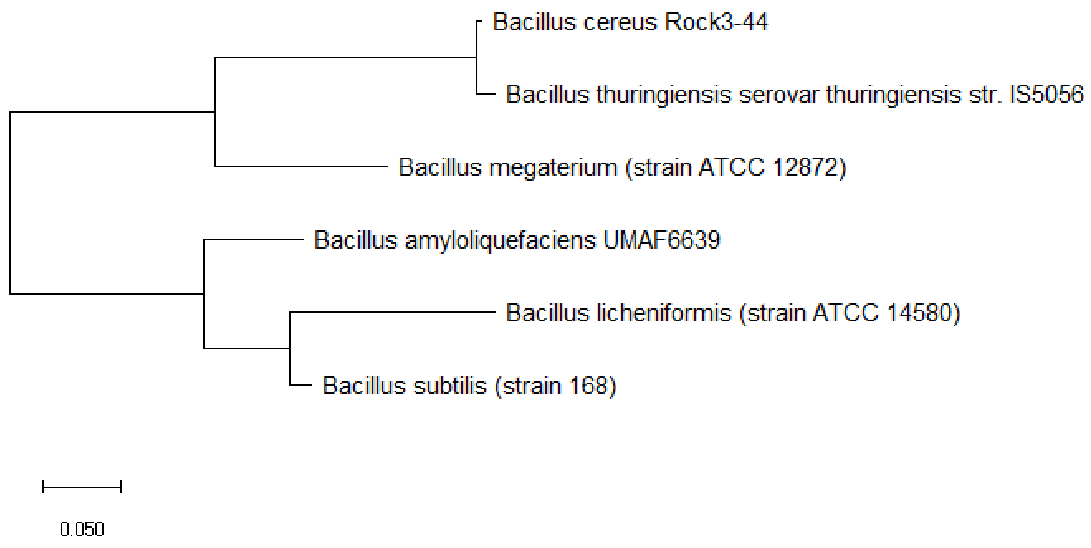


**Figure S22:** Maximum likelihood phylogenetic tree based on ribosomal protein L31 Type B of different *Bacillus* species.

Figure S22 shows the maximum likelihood phylogenetic tree based on ribosomal protein L31 Type B of different *Bacillus* species. The data revealed that ribosomal protein L31 Type B placed the phylogenetic position of *Bacillus licheniformis* wrongly as compared to that in the 16S rRNA phylogeny. Thus, ribosomal protein L31 Type B holds partial phylogenetic significance for describing the phylogeny of different *Bacillus* species.

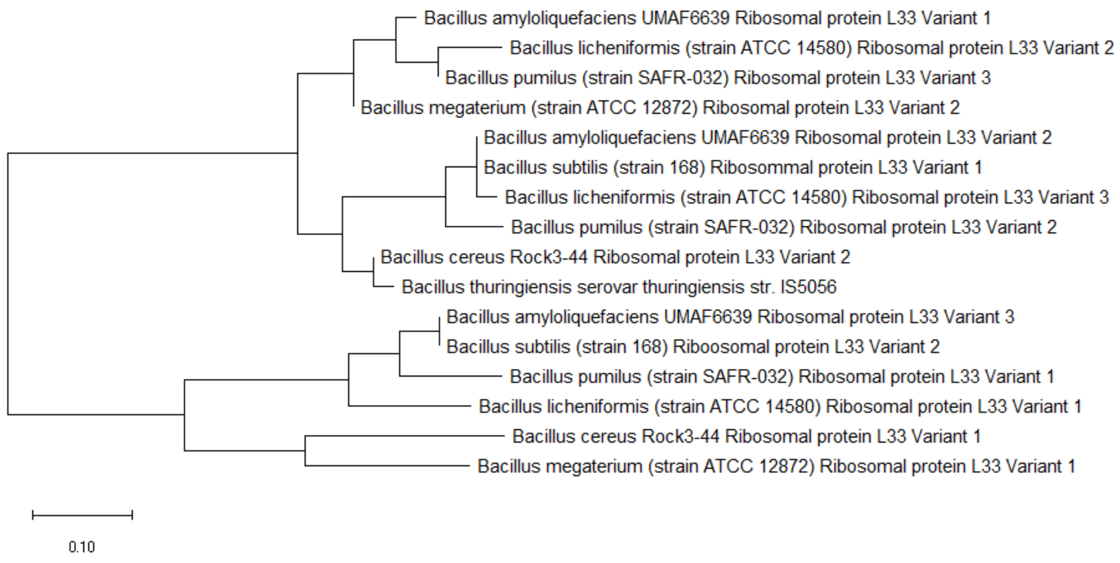


**Figure S23:** Maximum likelihood phylogenetic tree based on ribosomal protein L33 of different *Bacillus* species.

Figure S23 shows the maximum likelihood phylogenetic tree based on ribosomal protein L33 of different *Bacillus* species. As can be seen from Figure S23, there are many variants of ribosomal protein L33. The data revealed that ribosomal protein L33 could partially reproduce the phylogenetic tree based on 16S rRNA, and thus, hold partial phylogenetic significance for understanding the evolutionary relationships between different *Bacillus* species.

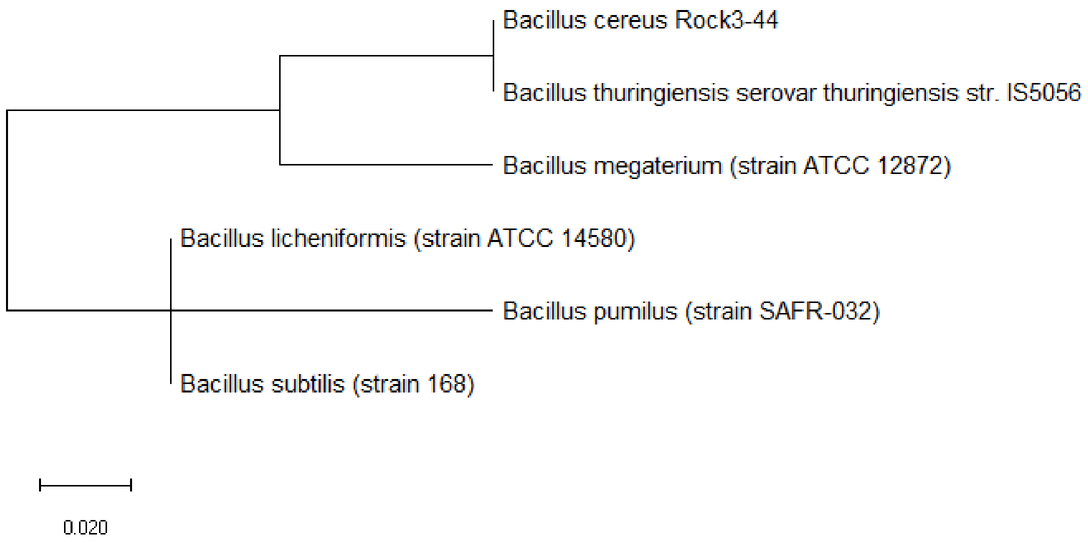

**Figure S24:** Maximum likelihood phylogenetic tree based on ribosomal protein L34 of different *Bacillus* species.

Figure S24 shows the maximum likelihood phylogenetic tree based on ribosomal protein L34 of different *Bacillus* species. The data revealed that ribosomal protein L34 could not reproduce one branch of the 16S rRNA phylogenetic tree, and thus, it does not hold phylogenetic significance for chronicling the evolutionary relationships between different *Bacillus* species.

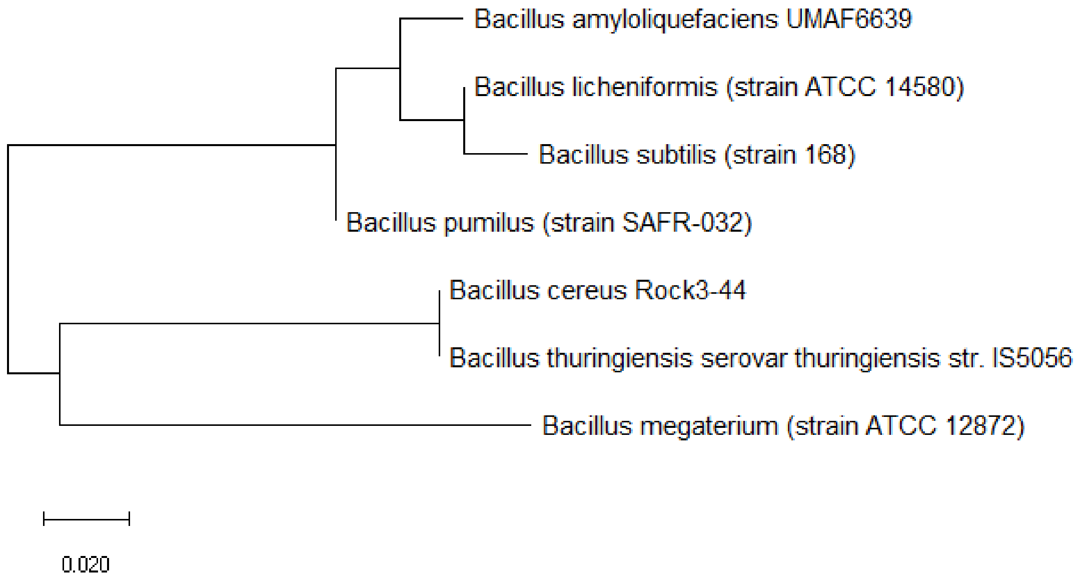


**Figure S25:** Maximum likelihood phylogenetic tree based on ribosomal protein L35 of different *Bacillus* species.

Figure S25 shows the maximum likelihood phylogenetic tree based on ribosomal protein L35 of different *Bacillus* species. The data revealed that apart from the misplacement of the phylogenetic position of *Bacillus subtilis* and *Bacillus pumilus*, ribosomal protein L35 could reproduce major branches of the 16S rRNA phylogenetic tree. Thus, ribosomal protein L35 holds partial phylogenetic significance for understanding the phylogeny of different *Bacillus* species.

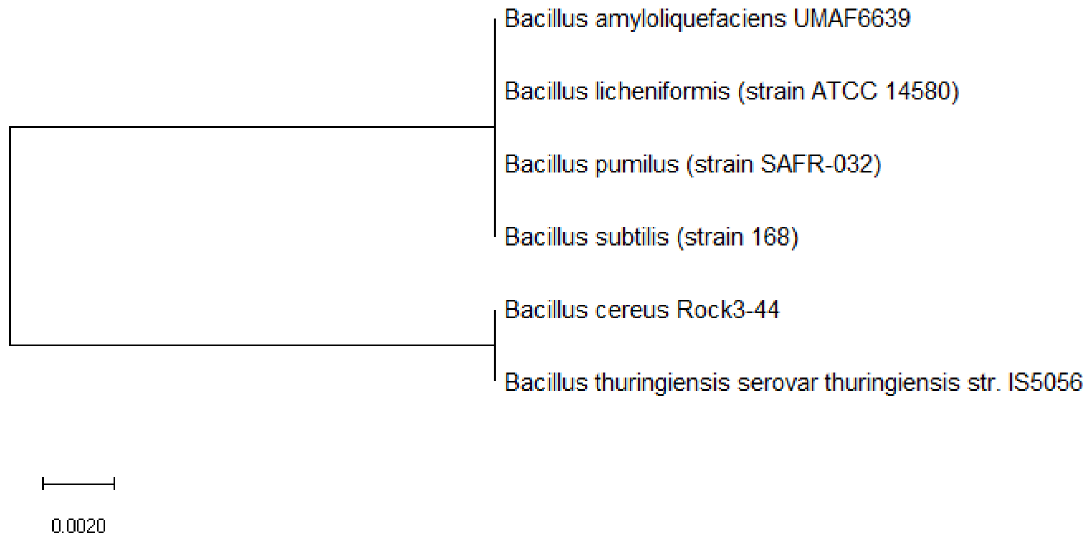


**Figure S26:** Maximum likelihood phylogenetic tree based on ribosomal protein L36 of different *Bacillus* species.

Figure S26 shows the maximum likelihood phylogenetic tree based on ribosomal protein L36 of different *Bacillus* species. The data revealed that the ribosomal protein L36 amino acid sequence is too highly conserved between different *Bacillus* species for it to explain the evolutionary relationships between the species. Thus, ribosomal protein L36 does not hold phylogenetic significance for this set of *Bacillus* species.

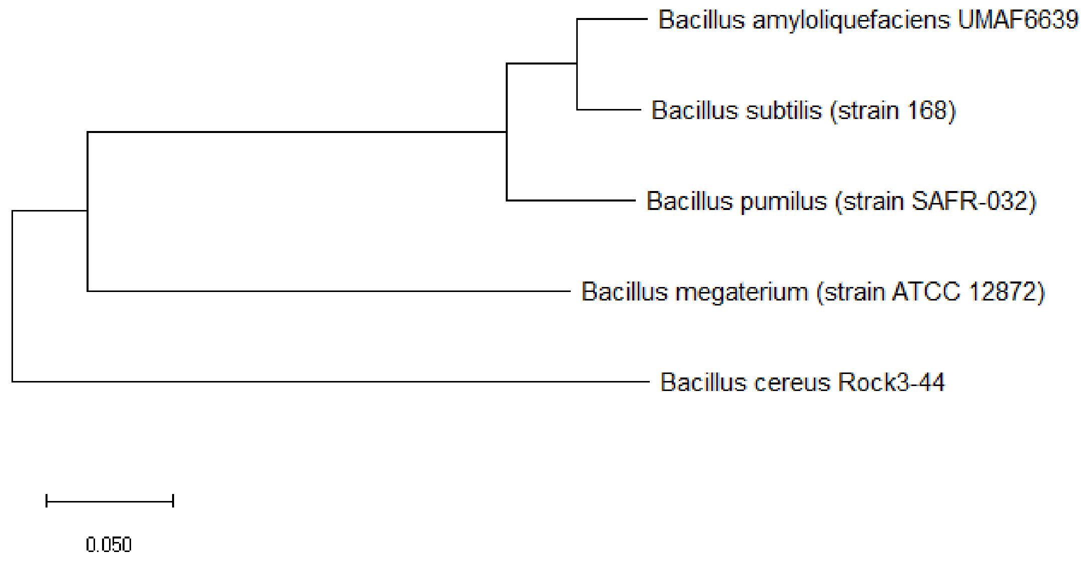

**Figure S27:** Maximum likelihood phylogenetic tree based on ribosomal protein S1 of different *Bacillus* species.

Figure S27 shows the maximum likelihood phylogenetic tree based on ribosomal protein S1 of different *Bacillus* species. The data revealed that ribosomal protein S1 could reproduce major branches of the 16S rRNA phylogenetic tree, and thus, it holds partial phylogenetic significance for explaining the phylogeny of different *Bacillus* species.

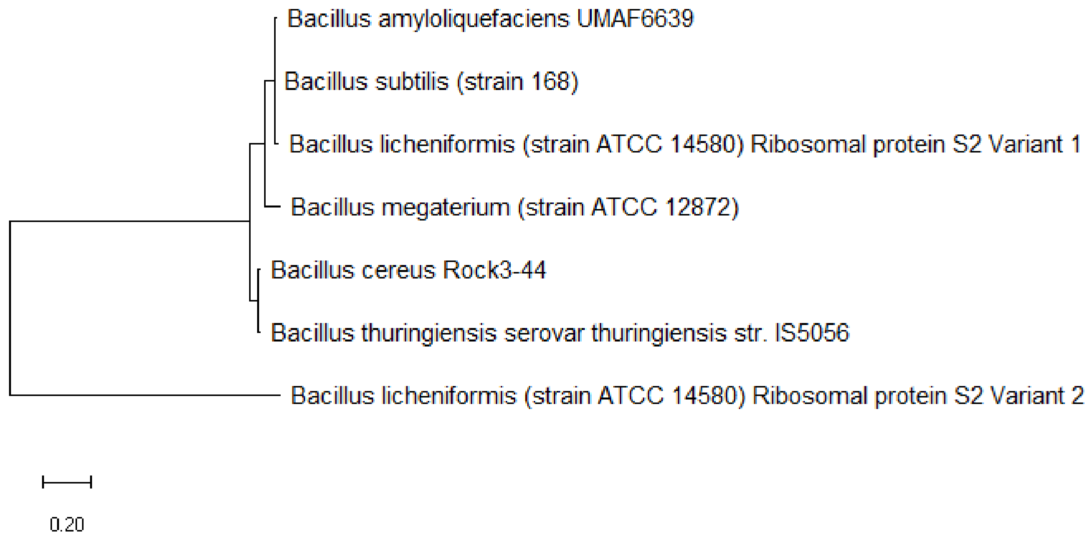


**Figure S28:** Maximum likelihood phylogenetic tree based on ribosomal protein S2 of different *Bacillus* species.

Figure S28 shows the maximum likelihood phylogenetic tree based on ribosomal protein S2 of different *Bacillus* species. The data revealed that the amino acid sequence of ribosomal protein S2 was too highly conserved to allow the chronicle of the evolutionary relationships between the different *Bacillus* species. Thus, ribosomal protein S2 does not hold phylogenetic significance.

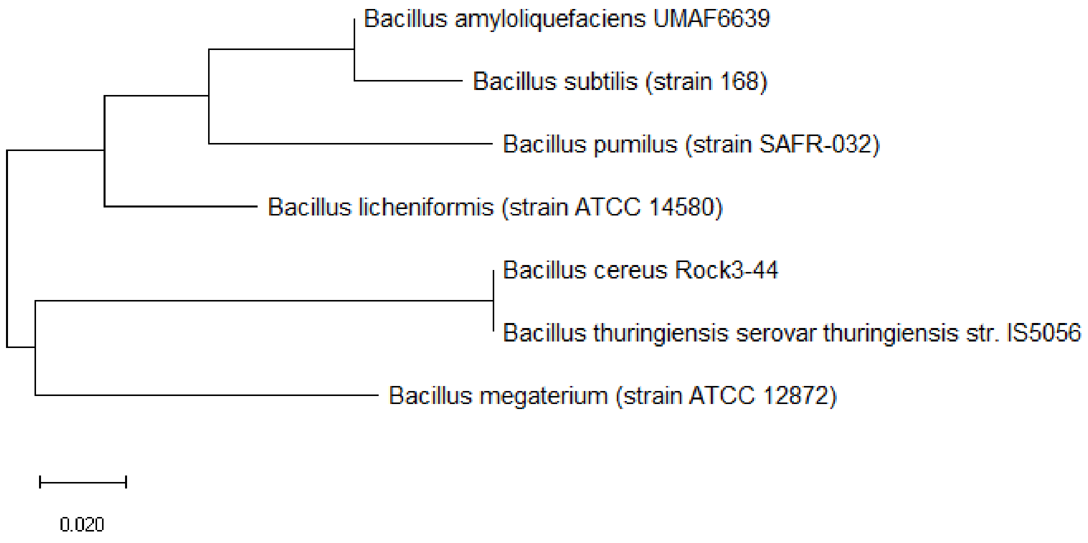


**Figure S29:** Maximum likelihood phylogenetic tree based on ribosomal protein S4 of different *Bacillus* species.

Figure S29 shows the maximum likelihood phylogenetic tree based on ribosomal protein S4 of different *Bacillus* species. The data revealed that ribosomal protein S4 could reproduce most of the branches of the 16S rRNA phylogenetic tree except for the placement of *Bacillus licheniformis* and *Bacillus pumilus*. Thus, ribosomal protein S4 holds partial phylogenetic significance for explaining the phylogeny of the different *Bacillus* species.

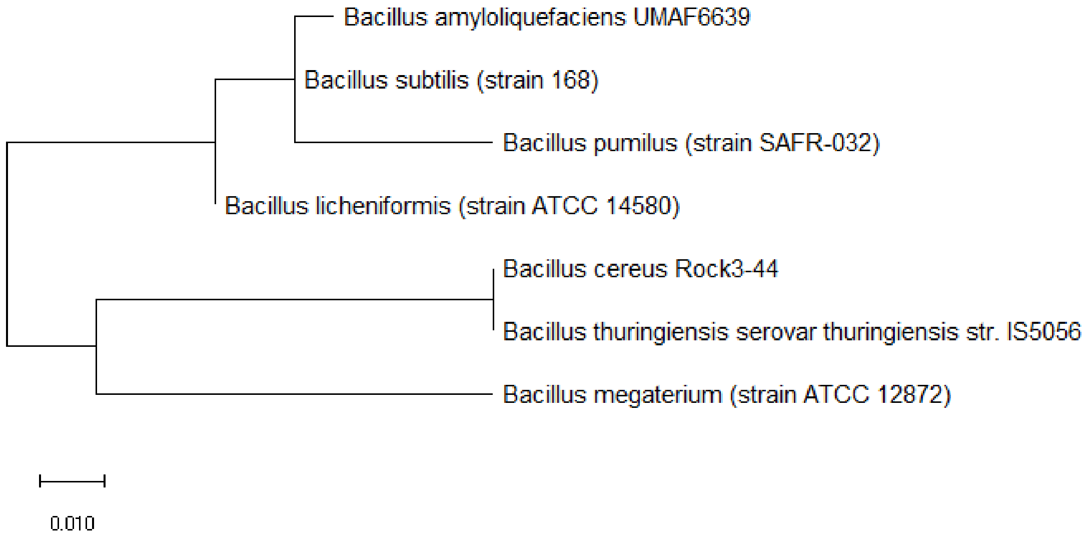


**Figure S30:** Maximum likelihood phylogenetic tree based on ribosomal protein S5 of different *Bacillus* species.

Figure S30 shows the maximum likelihood phylogenetic tree based on ribosomal protein S5 of different *Bacillus* species. The data revealed that except for *Bacillus licheniformis* and *Bacillus pumilus*, ribosomal protein S5 could place the other *Bacillus* species at the correct phylogenetic position relative to the phylogeny described by 16S rRNA. Thus, ribosomal protein S5 holds partial phylogenetic significance for explaining the phylogeny of different *Bacillus* species.

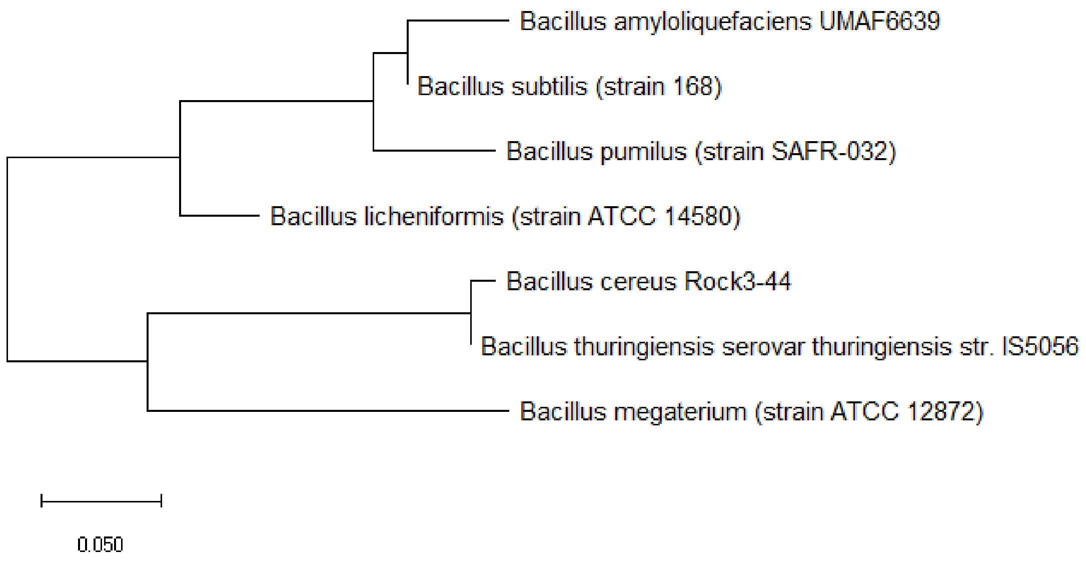


**Figure S31:** Maximum likelihood phylogenetic tree based on ribosomal protein S6 of different *Bacillus* species.

Figure S31 shows the maximum likelihood phylogenetic tree based on ribosomal protein S6 of different *Bacillus* species. The data revealed that ribosomal protein S6 could place most of the *Bacillus* species at the correct phylogenetic position relative to the phylogeny described by 16S rRNA. However, the phylogenetic position for *Bacillus licheniformis* and *Bacillus pumilus* were wrong. Thus, ribosomal protein S6 holds partial phylogenetic significance for explaining the evolutionary relationship between different *Bacillus* species.

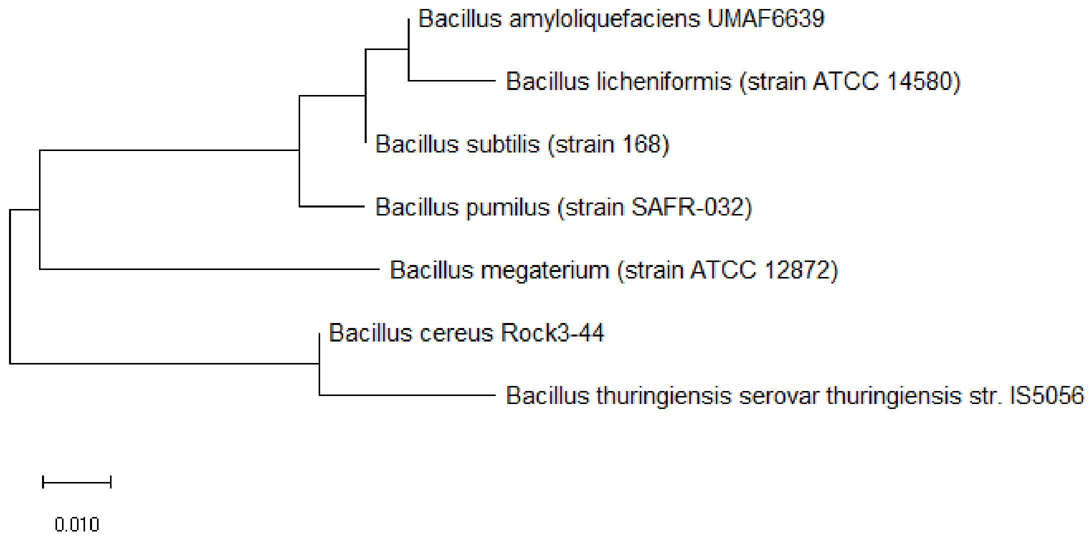


**Figure S32:** Maximum likelihood phylogenetic tree based on ribosomal protein S7 of different *Bacillus* species.

Figure S32 shows the maximum likelihood phylogenetic tree based on ribosomal protein S7 of different *Bacillus* species. The data revealed that ribosomal protein S7 could not place the phylogenetic position of *Bacillus subtilis* and *Bacillus licheniformis* correctly relative to the phylogeny described by 16S rRNA. Thus, ribosomal protein S7 holds partial phylogenetic significance for explaining the evolutionary relationships between different *Bacillus* species.

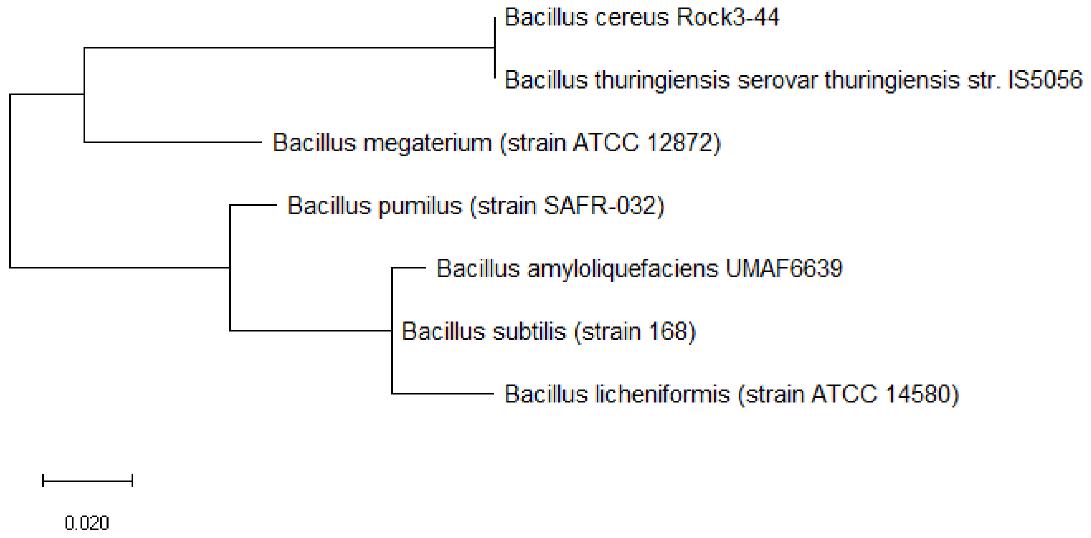

**Figure S33:** Maximum likelihood phylogenetic tree based on ribosomal protein S8 of different *Bacillus* species.

Figure S33 shows the maximum likelihood phylogenetic tree based on ribosomal protein S8 of different *Bacillus* species. The data revealed that ribosomal protein S8 could reproduce major branches of the 16S rRNA phylogenetic tree except for the wrong placement of *Bacillus licheniformis's* phylogenetic position. Thus, ribosomal protein S8 holds partial phylogenetic significance for explaining the phylogeny of different *Bacillus* species.

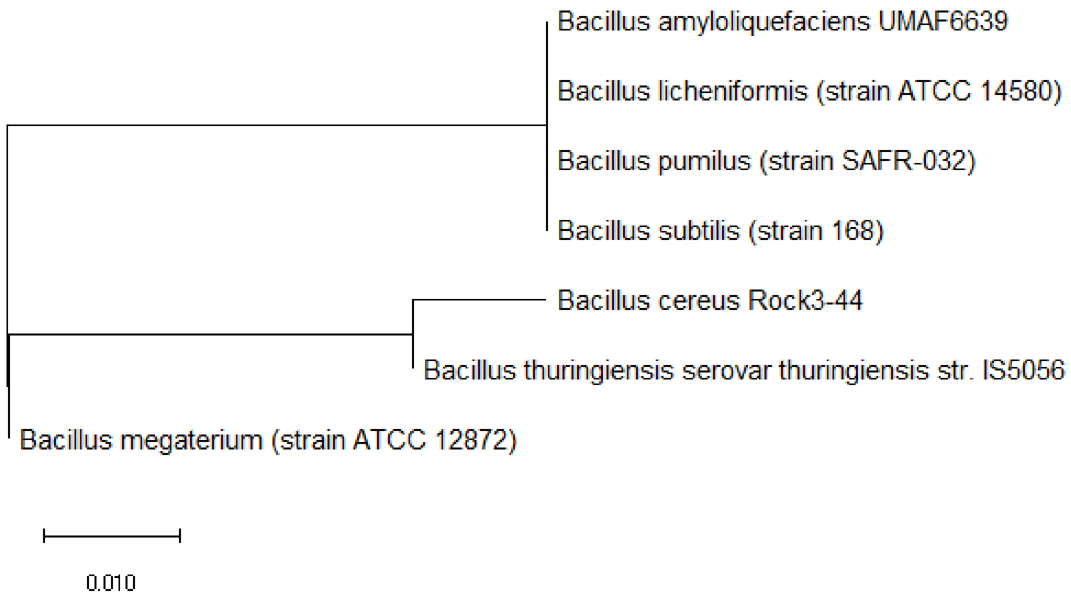


**Figure S34:** Maximum likelihood phylogenetic tree based on ribosomal protein S10 of different *Bacillus* species.

Figure S34 shows the maximum likelihood phylogenetic tree based on ribosomal protein S10 of different *Bacillus* species. The data revealed that ribosomal protein S10 was too highly conserved in amino acid sequence to encode sufficient sequence diversity to chronicle the evolutionary relatedness between different *Bacillus* species. Thus, ribosomal protein S10 does not hold phylogenetic significance.

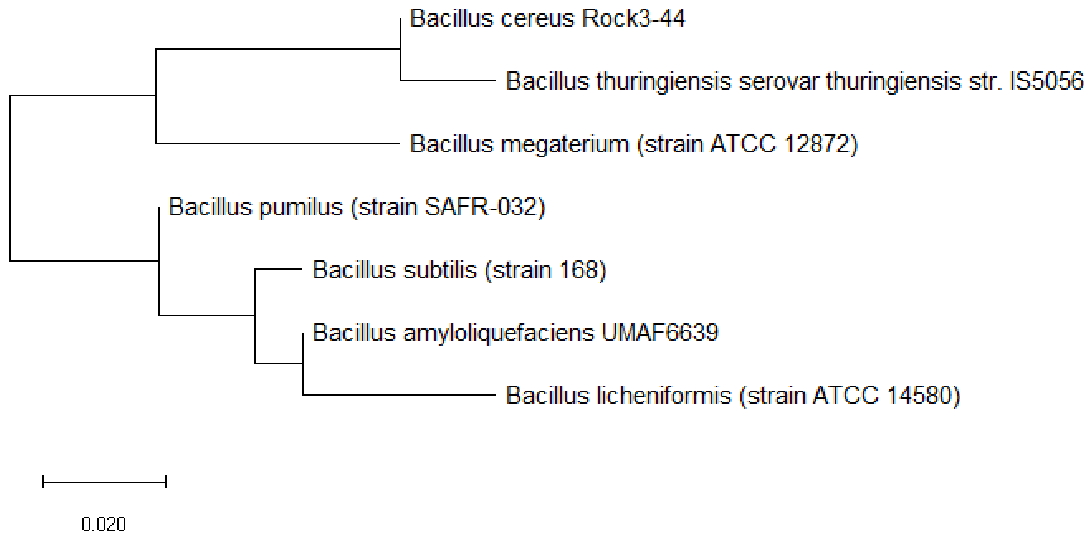


**Figure S35:** Maximum likelihood phylogenetic tree based on ribosomal protein S11 of different *Bacillus* species.

Figure S35 shows the maximum likelihood phylogenetic tree based on ribosomal protein S11 of different *Bacillus* species. The data revealed that except for the wrong placement of *Bacillus licheniformis's* phylogenetic position, ribosomal protein S11 could reproduce the major branches of the 16S rRNA phylogenetic tree. Thus, ribosomal protein S11 holds partial phylogenetic significance for explaining the phylogeny of different *Bacillus* species.

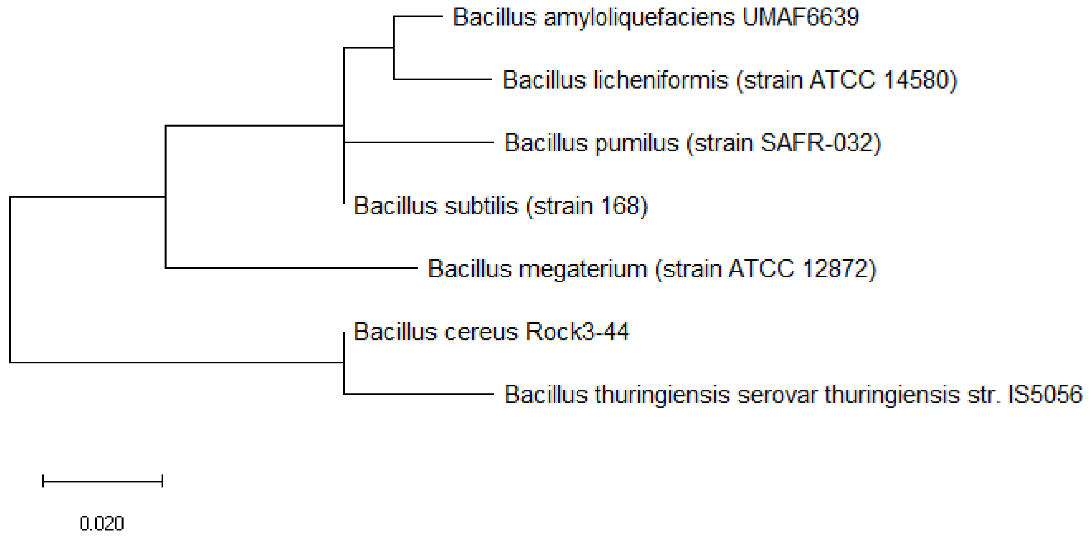


**Figure S36:** Maximum likelihood phylogenetic tree based on ribosomal protein S13 of different *Bacillus* species.

Figure S36 shows the maximum likelihood phylogenetic tree based on ribosomal protein S13 of different *Bacillus* species. The data revealed that except for *Bacillus subtilis*, ribosomal protein S13 could reproduce the major branches of the 16S rRNA phylogenetic tree for the set of *Bacillus* species. Thus, ribosomal protein S13 holds partial phylogenetic significance for explaining the phylogeny of the different *Bacillus* species.

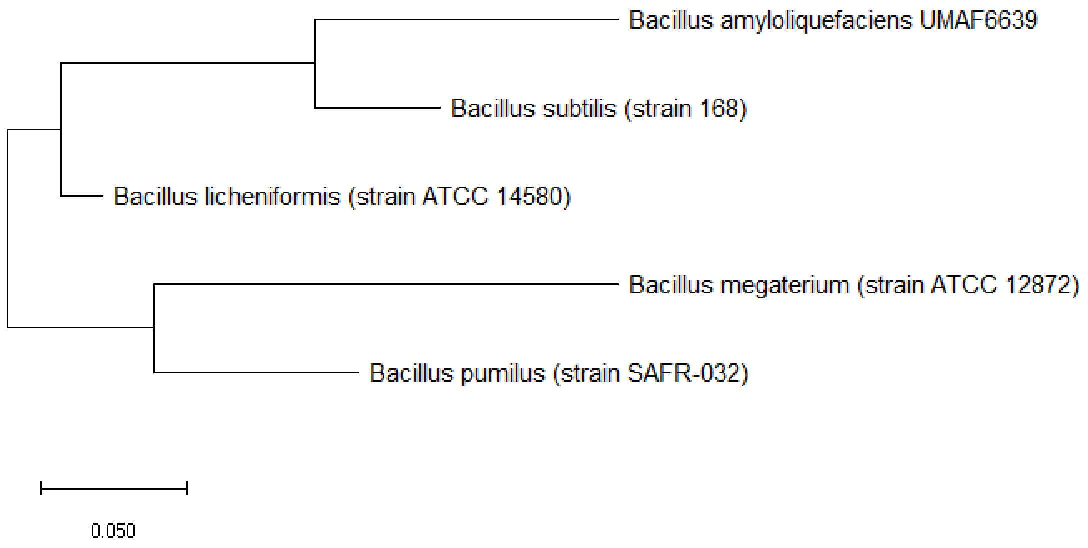


**Figure S37:** Maximum likelihood phylogenetic tree based on ribosomal protein S14 of different *Bacillus* species.

Figure S37 shows the maximum likelihood phylogenetic tree based on ribosomal protein S14 of different *Bacillus* species. As there are too few *Bacillus* species for meaningful comparison of the phylogenetic tree with that based on 16S rRNA, no conclusion could be drawn on whether ribosomal protein S14 hold phylogenetic significance for the different *Bacillus* species investigated.

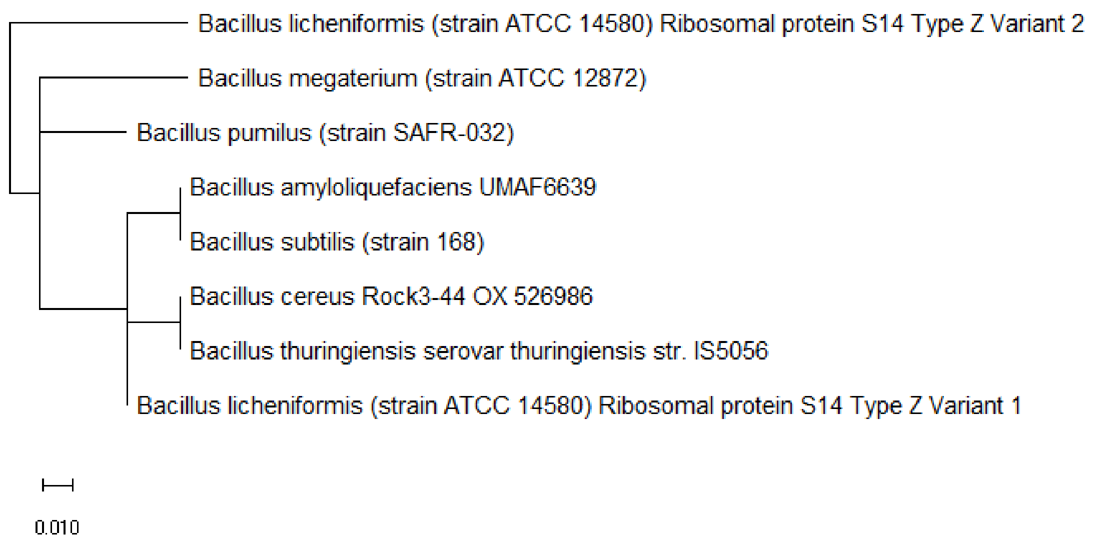


**Figure S38:** Maximum likelihood phylogenetic tree based on ribosomal protein S14 Type Z of different *Bacillus* species.

Figure S38 shows the maximum likelihood phylogenetic tree based on ribosomal protein S14 Type Z of different *Bacillus* species. The data revealed that there was no concordance between the phylogenetic tree of ribosomal protein S14 Type Z and that of 16S rRNA. Thus, ribosomal protein S14 Type Z does not hold phylogenetic significance for understanding the evolutionary trajectory of different *Bacillus* species.

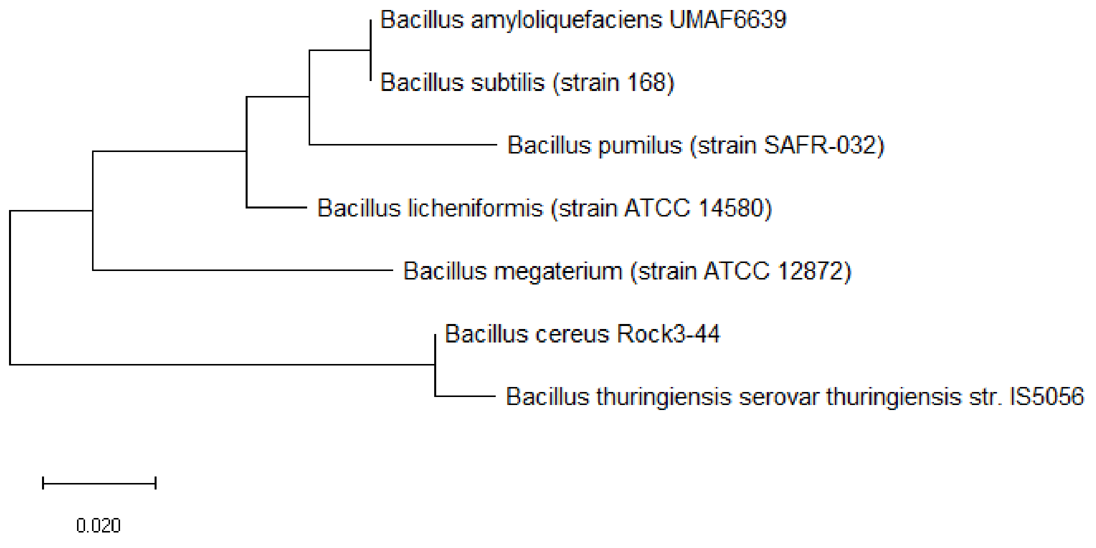


**Figure S39:** Maximum likelihood phylogenetic tree based on ribosomal protein S19 of different *Bacillus* species.

Figure S39 shows the maximum likelihood phylogenetic tree based on ribosomal protein S19 of different *Bacillus* species. The data revealed that except for the phylogenetic position of *Bacillus licheniformis* and *Bacillus pumilus*, ribosomal protein S19 could reproduce the major branches of the 16S rRNA phylogenetic tree for the *Bacillus* species investigated. Thus, ribosomal protein S19 holds partial phylogenetic significance for explaining the phylogeny of the different *Bacillus* species.

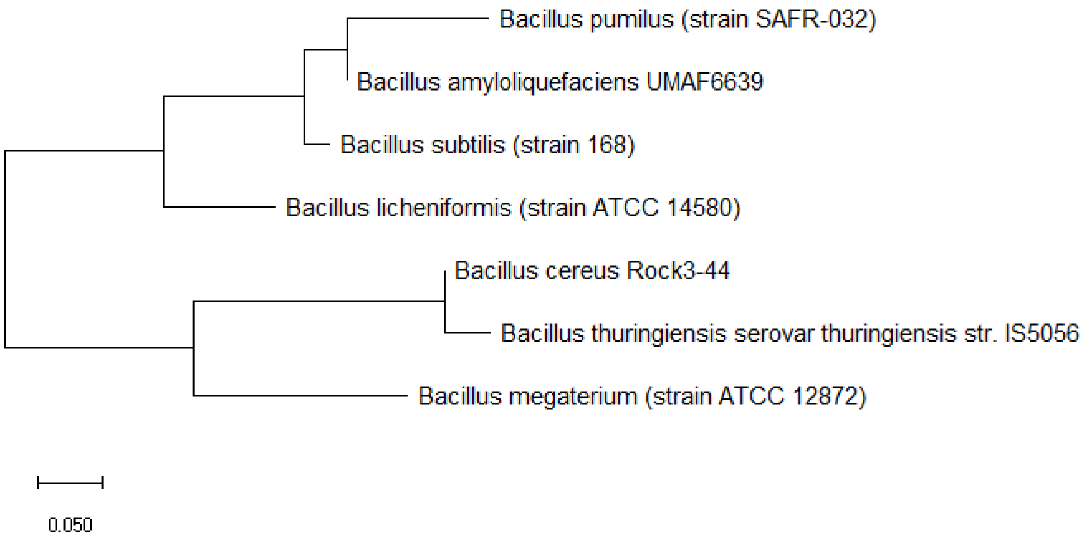


**Figure S40:** Maximum likelihood phylogenetic tree based on ribosomal protein S20 of different *Bacillus* species.

Figure S40 shows the maximum likelihood phylogenetic tree based on ribosomal protein S20 of different *Bacillus* species. The data revealed that ribosomal protein S20 could not explain the phylogeny of *Bacillus subtilis* and *Bacillus pumilus*, even though other branches of the 16S rRNA phylogenetic tree could be reproduced. Thus, ribosomal protein S20 holds partial phylogenetic significance for explaining the evolutionary relationships between different *Bacillus* species.

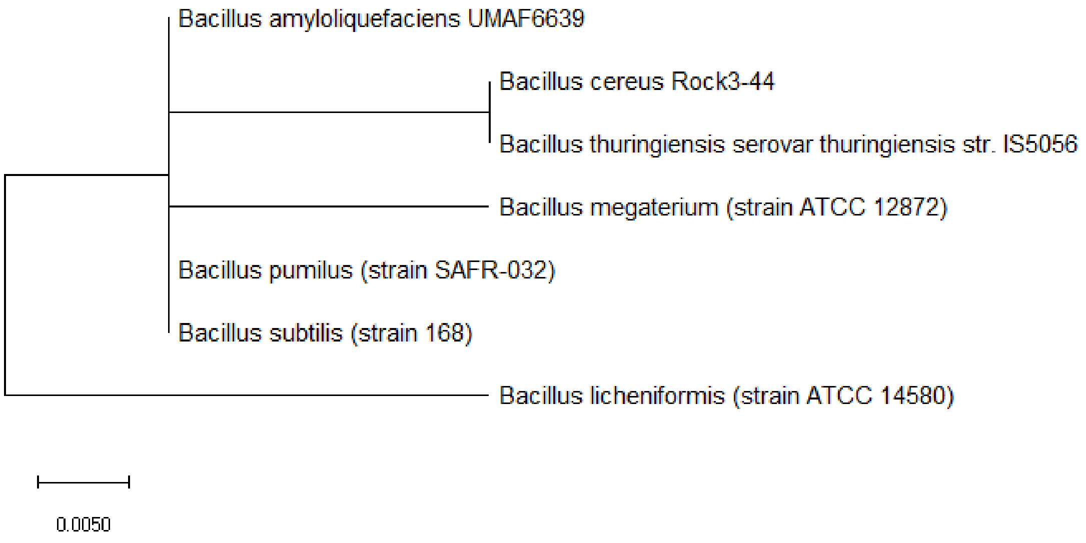


**Figure S41:** Maximum likelihood phylogenetic tree based on ribosomal protein S21 of different *Bacillus* species.

Figure S41 shows the maximum likelihood phylogenetic tree based on ribosomal protein S21 of different *Bacillus* species. The data revealed that ribosomal protein S21 lacks sequence diversity for chronicling the evolutionary history of the different *Bacillus* species given that the phylogenetic tree obtained revealed that the *Bacillus* species were too closely-related to each other. Thus, ribosomal protein S21 does not hold phylogenetic significance for explaining the phylogeny of different *Bacillus* species.

### Conflicts of interest

The author declares no conflicts of interest.

### Funding

No funding was used in this work.
